# Ultrasensitive fitness costs arise from membrane proteome displacement under protein overexpression

**DOI:** 10.1101/2025.02.12.637850

**Authors:** Janina Müller, Natawan Gadjisade, S. Andreas Angermayr, Gerrit Ansmann, Tobias Bollenbach

**Author notes:** CeMM Research Center for Molecular Medicine of the Austrian Academy of Sciences, 1090 Vienna, Austria (S. Andreas Angermayr).

## Abstract

Protein overexpression is crucial in many diseases and can lead to antibiotic resistance by increasing the expression of drug targets or resistance genes, such as membrane-localized efflux pumps. Using a high-throughput colony-imaging technique, we performed a quantitative genome-wide study of protein overexpression in *Escherichia coli*. Our results show that most membrane proteins impose steep overexpression costs, leading to an abrupt growth collapse beyond a critical expression threshold. By manipulating synthetic membrane proteins to target different translocation pathways, we ruled out saturation of these pathways as the cause of these fitness costs. Instead, we found that the growth collapse is driven by the displacement of endogenous membrane proteins, which we show directly using single-cell time-lapse imaging with fluorescently tagged membrane proteins in a microfluidic device. Displacement of as little as 10% of the membrane proteome trapped most bacteria in a non-growing state. Our findings enhance the quantitative understanding of expression costs, with implications for antibiotic resistance and the biotechnological production of membrane proteins.

## Introduction

Transcriptional regulation networks can ensure that proteins are produced at appropriate levels when needed, thereby minimizing the costs of unnecessary synthesis (Dekel and Alon, 2005; Keren et al., 2016). However, protein overexpression is a hallmark of many diseases, with broad biological and clinical implications (Sopko et al., 2006). A common way for bacteria to become resistant to antibiotics is to overexpress drug targets or resistance genes, such as efflux pumps, to counteract drug effects (Couce et al., 2012; Nikaido and Pagès, 2012; Palmer and Kishony, 2014; Soo et al., 2011). These resistance mechanisms incur fitness costs that determine the rate and trajectory of adaptive evolution under antibiotic pressure (Andersson, 2006). Ultimately, understanding the causes of protein overexpression costs is crucial for elucidating fundamental cellular constraints and their impact on adaptive responses to environmental stressors.

Quantitative studies of fitness costs of overexpression have largely focused on a few individual proteins, such as β-galactosidase (LacZ) (Eames and Kortemme, 2012; Stoebel et al., 2008) or mCherry (Kafri et al., 2015). In *Escherichia coli*, the fitness costs of overexpression have been meticulously quantified by expressing LacZ in the absence of its substrate or by expressing truncated, non-functional versions of the translation factor TufB, thereby eliminating any potential benefits of protein expression (Dong et al., 1995; Scott et al., 2010). When unnecessary proteins are overexpressed, they can occupy a large fraction of the proteome and effectively replace essential proteins due to the inherently limited protein synthesis capacity, thereby imposing a general fitness cost (Dong et al., 1995; Scott et al., 2010). This replacement also leads to increased cell size and DNA content by reducing the abundance of division-related proteins (Basan et al., 2015). In yeast and bacteria, quantitative analysis has revealed a simple linear relationship between fitness cost and the fraction of the proteome replaced by a gratuitous protein (Kafri et al., 2015; Scott et al., 2010), consistent with predictions from bacterial growth laws (Scott et al., 2010).

Beyond this general burden of overexpression, genome-wide overexpression studies have identified specific protein features that contribute to fitness costs. In yeast and other eukaryotes, proteins with high intrinsic disorder are particularly costly (Tomala and Korona, 2013; Vavouri et al., 2009), while in *E. coli*, high-cost proteins tend to be enriched in branched-chain amino acids (Chen et al., 2018). Across organisms, membrane proteins consistently impose particularly high costs (Chen et al., 2018; Gubellini et al., 2011; Tomala and Korona, 2013). However, due to methodological limitations, most genome-wide studies have used all-or-nothing overexpression or relied on crude fitness metrics such as colony formation defects, which primarily capture severe growth inhibition but fail to quantify more subtle fitness costs (Kitagawa et al., 2006; Sopko et al., 2006). More quantitative, high-resolution studies of genome-wide protein overexpression are needed to gain a deeper understanding of how protein features such as function, localization, and structure determine fitness costs.

Understanding the fitness costs of protein overexpression is also essential for the production of purified proteins for structural biology and biotechnology applications. High protein yields and minimized cellular fitness costs are interrelated goals. Membrane proteins, which are critical for signaling, transport, and as drug targets, are notoriously difficult to overexpress (Geertsma et al., 2008; Wagner et al., 2008), in part due to their high fitness costs. We focus on plasma membrane proteins (integral membrane proteins), a subset of membrane proteins that are integrated into the cytoplasmic membrane, and refer to them throughout as “membrane proteins” for simplicity. In *E. coli*, which serves as a key model system for studying membrane protein translocation, most membrane proteins are translocated or inserted into the membrane via the Sec pathway and rely on its core component, the SecYEG translocon (Crane and Randall, 2017; Kuhn et al., 2017). The SecYEG translocon is a highly conserved protein complex across organisms and is widely regarded as the main bottleneck limiting membrane protein expression (Wagner et al., 2007). Membrane proteins are targeted to the SecYEG translocon by the signal recognition particle (SRP), which recognizes the protein’s N-terminal signal sequences and localizes translating ribosomes to the plasma membrane. While some small membrane proteins can be integrated into the membrane without the translocation machinery or with the help of the insertase YidC alone, others require a varying number of components of the SecYEG translocon (Kuhn et al., 2017). To optimize protein yields, it is essential to identify whether the SecYEG translocon or other factors limit protein expression.

In this study, we employed a high-throughput colony-imaging technique to quantify genome-wide overexpression costs in *E. coli* by measuring microbial growth rates with high precision. Our analysis confirmed that membrane proteins impose exceptionally high fitness costs, but also revealed a surprising critical threshold at which membrane protein overexpression triggers an abrupt and irreversible growth collapse. Through experiments with a series of synthetic membrane proteins of varying hydrophilicity, fluorescently tagged membrane proteins, single-cell time-lapse imaging and microfluidics, we show that this collapse is driven by the competitive displacement of endogenous membrane proteins rather than by saturation of the SecYEG translocon. This displacement traps bacteria in a dysfunctional membrane proteome state from which they rarely recover. Our findings advance our quantitative understanding of protein expression costs, provide insights into the fitness burden of efflux-mediated antibiotic resistance, and inform strategies to maximize membrane protein yields in biotechnology.

## Results

### Quantifying genome-wide fitness costs of protein overexpression

To enable comprehensive genome-wide measurements of the fitness costs associated with protein overexpression, we developed a high-throughput method for bacterial growth analysis using colony time-lapse imaging. Briefly, we arrayed up to 1,536 *E. coli* colonies on agar plates and imaged their growth over 26 hours using a colony processing robot (Methods). We then obtained growth curves from the background-corrected mean pixel intensity of the inner portion of each colony (Figure 1a–c), which correlates strongly with colony biomass and reliably captures early exponential growth over approximately two orders of magnitude (Supplementary Figure 1). Focusing on the early growth phase minimizes interference from neighboring colonies and edge effects (Supplementary Figure 2), which become a substantial problem at later stages (Baryshnikova et al., 2010). Batch effects were also modest (Supplementary Figure 2). This method provides growth measurements of comparable quality to optical density readings in liquid culture (Supplementary Figure 3). However, unlike liquid culture methods, which are typically limited to 96 cultures per plate due to the need for vigorous shaking, this time-lapse colony imaging approach offers higher throughput: a basic imaging robot can measure over 65,000 growth curves in a single run (Figure 1a; Methods). This technique thus enables fine-grained, systematic studies of genome-wide strain libraries without sacrificing data quality.

**Figure 1:**
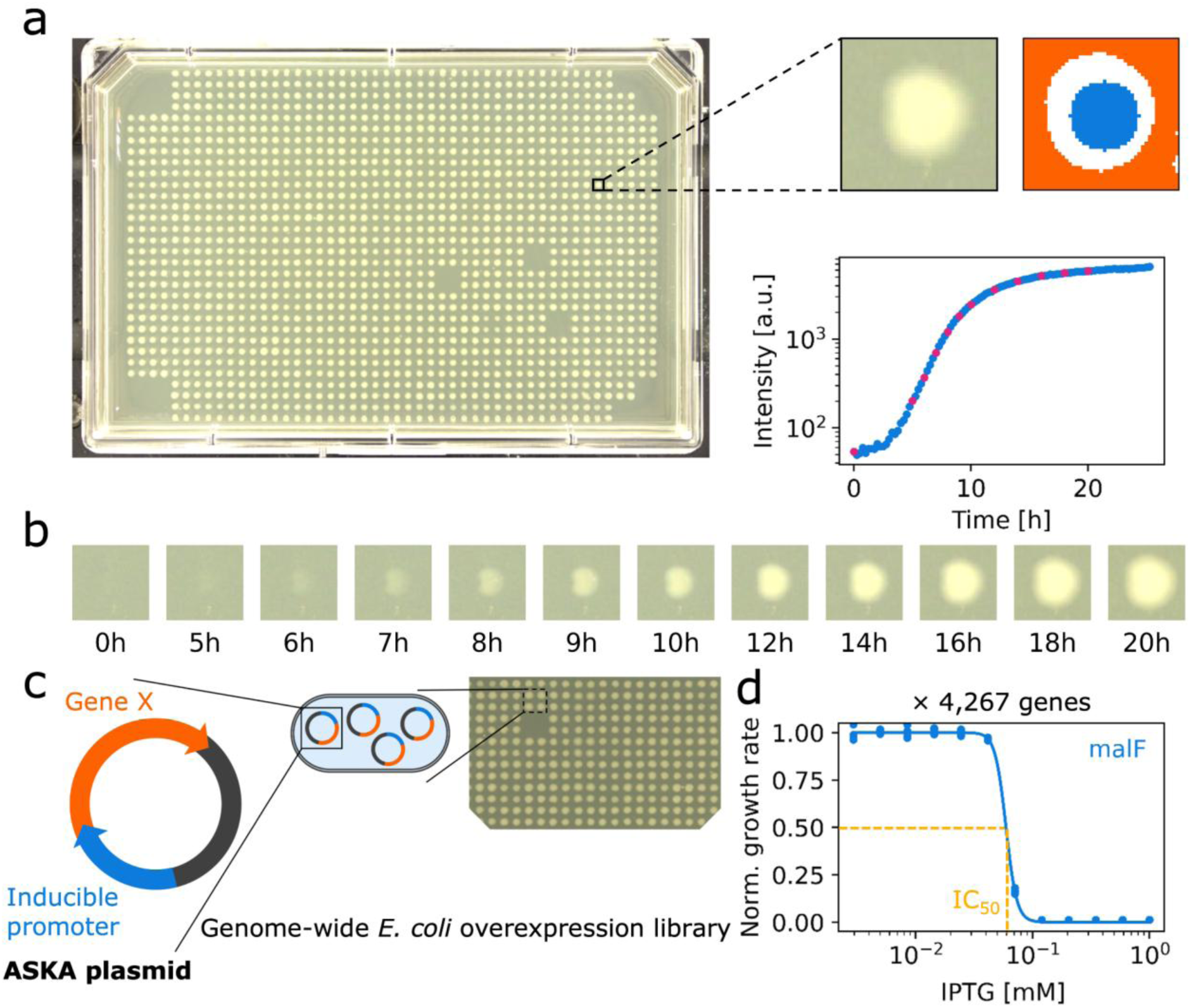
High-throughput quantification of overexpression dose-response curves using bacterial colony imaging. **a)** Example plate showing 1,536 colonies after 20 hours of growth (left panel). For each colony, growth curves (bottom right) are generated by subtracting the mean background intensity (orange) from the mean intensity of the colony center (blue) at each time point. The magenta points in the growth curve correspond to the images shown in panel b). **b)** Time series of a single colony from a plate containing 1,536 colonies. The intensity of the center of the colony increases before radial growth of the colony begins. The images show the colony immediately after pinning (0 hours) and at later time points. **c)** Schematic of the *E. coli* overexpression library used in the experiments. Each strain contains a high-copy-number plasmid with an inducible promoter (Kitagawa et al., 2006). **d)** Example overexpression dose-response curve for *malF* (blue dots). All 4,267 dose-response curves were fitted with a Hill function (blue line) to obtain the IC_50 (_dashed yellow line) and the Hill coefficient *n*, which measures its steepness.

Using this technique, we systematically characterized the fitness costs of continuously varying levels of protein overexpression across the *E. coli* genome. We measured growth rates for all 4,267 strains in the ASKA overexpression library (Figure 1d) (Kitagawa et al., 2006) at finely resolved, exponentially spaced inducer (IPTG) concentrations in quadruplicate (Methods). This generated an overexpression dose-response curve for each gene, characterizing the fitness cost as a function of inducer concentration (Figure 1d). Experiments with translational GFP fusions showed that IPTG concentrations are a reliable proxy for protein levels within the relevant experimental range (Supplementary Figure 4), consistent with findings that protein levels in bacteria are predominantly determined at the promoter level (Balakrishnan et al., 2022). The expression level of individual proteins at a given inducer concentration may vary due to differences in secondary mRNA structures that influence ribosome accessibility and codon usage, among other factors. However, we are primarily interested in comparing larger groups of proteins, where these differences are expected to average out. Therefore, we used the inducer concentration that reduces growth by 50% (IC_50)_ as a quantitative measure of overexpression cost for each gene (Figure 1d). Notably, the overexpression dose-response curve also reveals how sensitive the fitness is to changes in expression level, quantified by the Hill coefficient *n*, which measures the curve’s steepness around the IC_50_ (Figure 1d; hereafter, the word *sensitive* always refers to this quantity). Together, this dataset (Supplementary Table 1) provides a genome-wide quantitative characterization of the fitness cost of overexpression and its sensitivity to changes in expression level.

By systematically analyzing this dataset, we found key protein features that strongly influence the cost of overexpression and its sensitivity. While each protein has its own function, making overexpression costs idiosyncratic, we can identify the role of specific protein properties by analyzing larger groups of hundreds of genes. Gene ontology (GO) enrichment analysis showed that membrane proteins incur drastically higher fitness costs compared to cytosolic proteins (*p* < 10^−95^, Mann–Whitney *U* test; Figure 2a and b), consistent with previous findings (Chen et al., 2018; Gubellini et al., 2011; Osterberg et al., 2006; Tomala and Korona, 2013). Most of the 1,095 membrane proteins in our dataset have an IC_50 o_f about 0.08 mM, reflecting high fitness costs, while the 1,387 cytosolic proteins tend to have higher IC_50,_ 1 mM and higher, reflecting lower fitness costs. We validated this result by directly quantifying expression levels using 78 protein-GFP fusions (Supplementary Figure 5). We also observed high overexpression costs for the 424 DNA-binding proteins, including transcription factors (Figure 2a, Supplementary Figure 6a). Note that the IC_50 o_f many membrane and DNA-binding proteins is around 0.08 mM (Figure 2b), partly because expression levels increase steeply at this concentration of IPTG (Supplementary Figure 7c). In contrast, certain enzymes, including oxidoreductases, isomerases and lyases, as well as metal-ion-binding proteins, had particularly low costs (Supplementary Figure 6b). Protein length showed a weak but highly significant correlation with expression cost (Spearman rank correlation *r* = −0.24, *p* < 10^−44^), reflecting higher fitness costs of larger proteins, as expected (Figure 2c).

**Figure 2:**
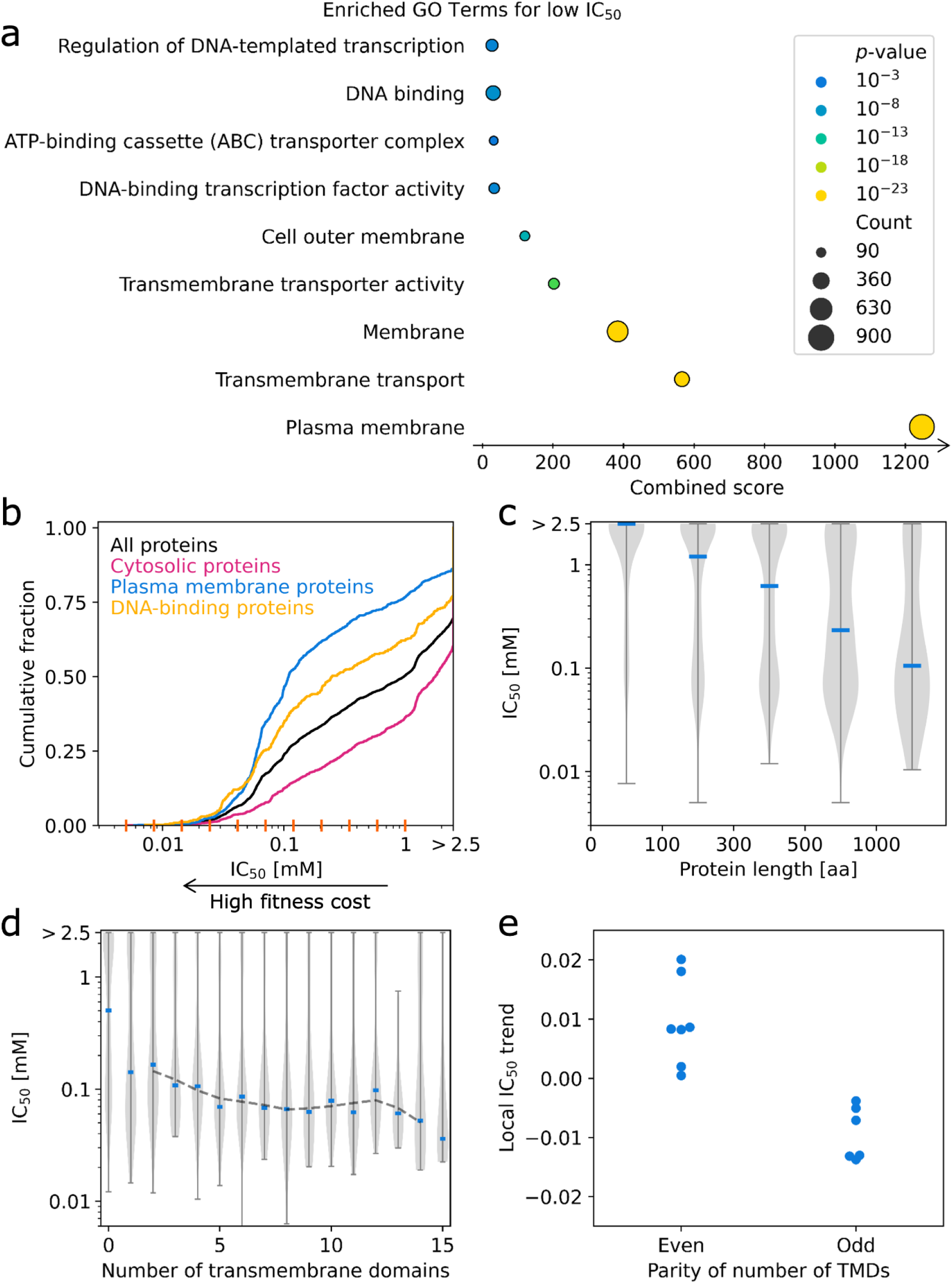
Membrane proteins have extremely high overexpression costs, which increase with the number of transmembrane domains (TMDs). **a)** Gene ontology enrichment analysis for proteins with low IC_50 v_alues (IC_50 <_ 0.5 mM) and with *p*-values below 10^−3^. **b)** Empirical cumulative distribution functions of IC_50 f_or all proteins (black) and by category (colors) (Methods). The orange markers on the x-axis indicate the IPTG concentrations sampled in the experiment. The IC_50 f_or many proteins exceeds the highest IPTG concentration used (2.5 mM). Therefore, the distribution functions end below one on the far right end (>2.5 mM); the remaining proteins have a higher IC_50,_ which cannot be determined from our data. Increased expression costs of membrane proteins are also observed as a function of the measured expression level (Supplementary Figure 5). **c)** Violin plot of IC_50 f_or different protein lengths. Median IC_50 v_alues (blue markers) decrease with increasing protein length, indicating higher fitness costs for longer proteins. **d)** Violin plot of IC_50 v_alues for membrane proteins versus number of TMDs. Blue markers indicate median IC_50 v_alues. An alternating pattern is observed, with a tendency for an odd number of TMDs to result in higher costs than an even number. The dashed gray line shows the running average ¼ x_n-1 +_ ½ x_n +_ ¼ x_n+1 o_ver the median IC_50 v_alues. **e)** Local IC_50 t_rend as the deviation from the running average for an odd or even number of TMDs. Membrane proteins with an odd number of TMDs tend to have lower IC_50 v_alues compared to the running average, while those with an even number of TMDs have higher IC_50 v_alues. This pattern was validated by bootstrapping of the data shown in panel **d** (Methods), confirming its occurrence in 97% of cases.

To elucidate the high fitness costs of membrane proteins, we examined additional protein characteristics related to the membrane area they occupy and the difficulty of their insertion. First, we found that the number of transmembrane domains (TMDs) correlates with expression costs (Spearman rank correlation r = −0.33, p < 10^-38^): proteins with more TMDs have higher expression costs (Figure 2d). Notably, proteins that localize to the membrane but lack TMDs – such as cytosolic proteins that are part of membrane-associated complexes – have much lower expression costs than those with one or more TMDs (Figure 2d). This suggests that membrane insertion is a key factor in the cost of membrane protein expression. Consistent with this view, we observed a trend that proteins with an odd number of TMDs tend to have higher expression costs than those with an even number (*p* = 0.03 from bootstrapping; Figure 2e). This might reflect the need for the terminal tail of a protein with an odd number of TMDs to cross the hydrophobic membrane, which is likely energetically unfavorable due to the typically hydrophilic nature of these tails. Second, we found that protein hydrophobicity, calculated as the average of the hydrophobicity indices of its amino acid sequence (Methods), was weakly correlated with overexpression cost (Spearman rank correlation *r* = −0.21, p < 10^−33^; Supplementary Figure 8). This correlation may be driven by the fact that hydrophobicity serves as another proxy for occupied membrane space, since transmembrane domains are predominantly composed of hydrophobic residues. These data suggest that the fitness cost of membrane proteins is influenced by various factors, including their size, the number of TMDs, their hydrophobicity, and whether the protein is merely associated with membrane-localized complexes. This offers a plausible explanation for the observed variability in fitness costs of membrane protein overexpression (Figure 2b).

Membrane proteins are much more sensitive to changes in expression level than cytosolic proteins. Their overexpression dose-response curve is steep, with growth rates dropping abruptly to zero once a critical threshold is exceeded. This high sensitivity is reflected in a median Hill coefficient of 6.7 for membrane proteins, compared to only 2.7 for cytosolic proteins (*p* = 10^−12^, Mann–Whitney *U* test; Figure 3a and b). Plotting dose-response curves against directly measured expression levels confirmed increased sensitivity to membrane protein overexpression compared to that of cytosolic proteins (Supplementary Figure 5a). GO enrichment analysis showed that genes with high Hill coefficients were strongly enriched for membrane proteins and related annotations (Supplementary Figure 9). To assess growth rate as a function of proteome fraction rather than inducer concentration, we estimated proteome fractions for each inducer concentration by comparing the cost of *lacZ* overexpression to previous studies that measured this cost as a function of proteome fraction (Supplementary Figure 7 and 7; Methods). Since inducer concentration controls protein copy number, but proteome fraction is measured by mass, we adjusted these fractions to account for the length of each individual protein relative to LacZ. Using this estimation, membrane proteins showed a sharply concave relationship, with growth dropping abruptly to zero around a proteome fraction of about 3% (Supplementary Figure 10). In contrast, overexpression of gratuitous cytosolic proteins resulted in a gradual, linear decline in growth rate, approaching zero at a proteome fraction of about 50% (Supplementary Figure 10), as previously reported (Scott et al., 2010). These results suggest that even modest levels of membrane protein overexpression push the cell into a sensitive regime where any further increase leads to an abrupt collapse in growth.

**Figure 3:**
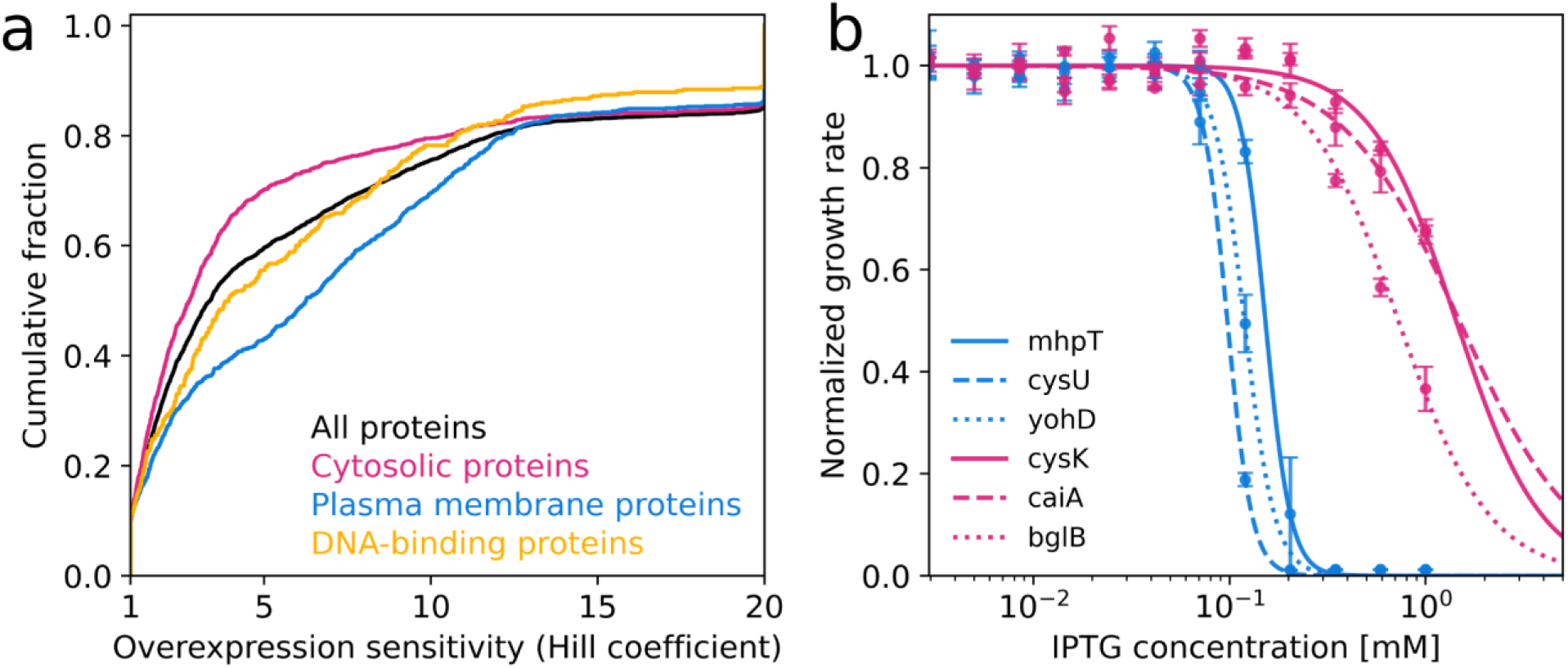
Bacterial growth declines sharply once a critical threshold of membrane protein expression is exceeded. **a)** Empirical cumulative distributions of Hill coefficients (Methods). Plasma membrane proteins (blue) typically have higher Hill coefficients than cytosolic proteins (magenta). DNA-binding proteins (yellow) also tend to have higher Hill coefficients, but to a lesser extent. **b)** Example dose-response curves for the membrane proteins MhpT (solid blue line), CysU (dashed blue line), and YohD (dotted blue line), and the cytosolic proteins CysK (solid magenta line), CaiA (dashed magenta line), and BglB (dotted magenta line). These curves represent approximately the median parameter values for membrane and cytosolic proteins across the genome: IC_50,MhpT =_ 0.15 mM, n_MhpT =_ 6.8, IC_50,CysU =_ 0.10 mM, n_CysU =_ 6.9, IC_50,YohD =_ 0.12 mM, n_YohD =_ 5.6, IC_50,CysK =_ 1.43 mM, n_CysK =_ 2.00, IC_50,CaiA =_ 1.49 mM, n_CysK =_ 1.5, and IC_50,BglB =_ 0.73 mM, n_BglB =_ 1.9. Dose-response data for all proteins are in Supplementary Table 1. A relatively sharp decline in growth rate for membrane proteins is also observed as a function of the measured expression level (Supplementary Figure 5).

The extreme cost and sensitivity of membrane protein overexpression is consistent with the view that there is a severely limited capacity for membrane proteins. This is plausible because the membrane is highly crowded, with up to 50% of its area occupied by proteins in *E. coli* (Kadner, R.J., 1996). At maximum induction, LacZ accounts for about 38% of the total proteome (by mass) in our experiments (Supplementary Figure 7). For an average membrane protein with two or more TMDs, which is considerably smaller than LacZ, the maximum achievable expression level in our experiments would correspond to 14% of the total proteome, or about half of the endogenous membrane proteome. However, growth collapses well before this point (Supplementary Figure 10), likely because essential membrane proteins, such as nutrient transporters, components of the electron transport chain, and ATP synthase, compete with the overexpressed protein for limited membrane space.

We hypothesized that membrane protein overexpression leads to the displacement of essential membrane proteins due to spatial constraints within the membrane, leading to a collapse of cell growth once a critical threshold of overexpression is exceeded (“displacement hypothesis”). An alternative explanation is that overexpression overloads the membrane-protein translocation pathway (Wagner et al., 2007). The displacement hypothesis is consistent with the observed correlations between overexpression cost and protein length (Figure 2c) and the number of TMDs (Figure 2d), which are close proxies for the occupied membrane area. At the same time, the overload on translocation pathways could also be exacerbated with increasing protein size, as larger proteins may occupy the translocation machinery for longer. While our observations show that this cost is not primarily due to an event that occurs only once per protein produced, such as translation initiation or targeting by SRP, they are consistent with both the displacement hypothesis and the possibility that an overload of the translocation pathway contributes to the effect.

### Load on the SecYEG translocon due to membrane-protein overexpression

To disentangle the effect of protein size on the cost of overexpression from the effect of load on the translocation pathway, we used the M13 procoat protein as a model membrane protein and varied its dependence on the Sec pathway (Figure 4a) (Hariharan et al., 2021). M13 procoat is the major coat protein (pVIII), i.e. the main component of the capsid, of the filamentous coliphage M13 (Rakonjac et al., 2011; Wickner and Killick, 1977). In the absence of other phage proteins and viral DNA, it serves as a gratuitous protein that integrates into the plasma membrane. Wild-type M13 procoat membrane insertion requires YidC but not the SecYEG translocon (Hariharan et al., 2021; Soman et al., 2014). These properties make it possible to modify the dependence on the Sec pathway while keeping the protein size constant, thus separating the specific contributions of size and translocation load from other costs of membrane protein overexpression.

**Figure 4:**
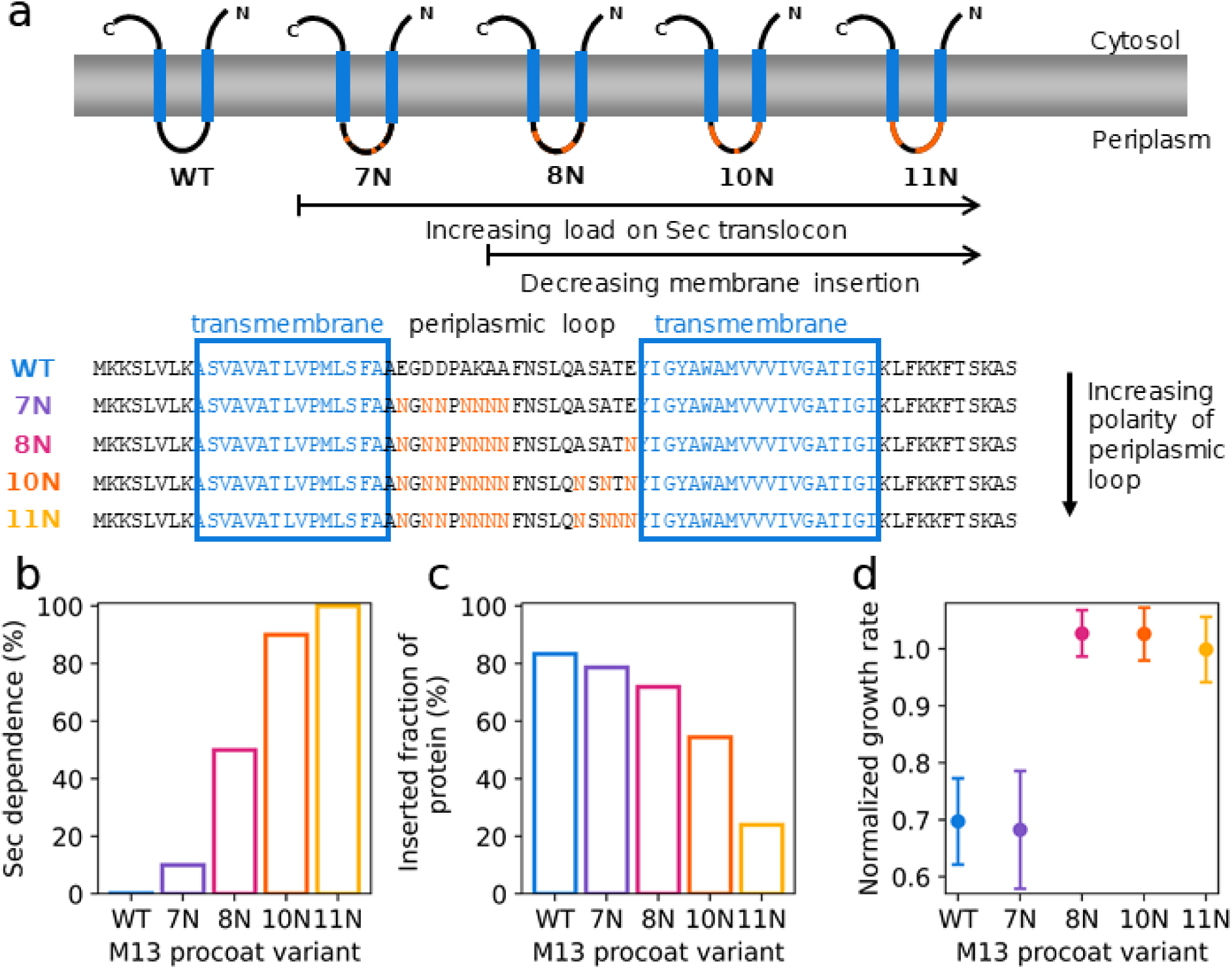
Increasing the load on the translocation pathway reduces the fitness cost of overexpression. **a)** Schematic of the M13 procoat protein mutant variants used in this study, adapted from (Hariharan et al., 2021b). The amino acid sequence of the periplasmic loop of this transmembrane protein was systematically modified by the addition of up to 11 hydrophilic arginine residues. **b)** Dependence of membrane insertion on the SecYEG translocon increases with additional hydrophilic residues in the periplasmic loop (from 7N to 11N) (Hariharan et al., 2021). **c)** The fraction of M13 procoat proteins successfully inserted into the membrane decreases with additional hydrophilic residues in the periplasmic loop (Methods).| **d)** Growth rate at maximum expression for each mutant. The cost of overexpression in the wildtype and 7N mutant is alleviated with increasing Sec-dependence and decreasing inserted fraction (8N,10N,11N).

To vary the dependence on the Sec pathway, we used five synthetic M13 procoat variants with increasing hydrophilicity of the periplasmic loop. Hydrophilicity is gradually increased by replacing an increasing number of amino acid residues with asparagine (N), a polar amino acid (Hariharan et al., 2021). As hydrophilicity increases, so does the dependence on the Sec pathway (Figure 4b). The variant with seven additional asparagine residues (7N) already requires the SecYEG translocon to be inserted with a similar probability as the WT protein, but it can still insert with lower efficiency in the absence of the SecYEG translocon; for the 8N and 10N variants, insertion efficiency decreases further and the inserted fraction of the protein is reduced compared to the WT and 7N variants; and the 11N variant largely fails to insert into the membrane, even with the help of the SecYEG translocon (Hariharan et al., 2021). Thus, in parallel with increasing dependence on the Sec pathway, increasing hydrophilicity reduces the propensity for membrane insertion (Figure 4c; Methods). Taken together, the series of synthetic membrane protein variants provides us with a means to increase the load on the Sec pathway and reduce the probability of successful membrane insertion, while maintaining the size and expression level of the protein.

Overexpression cost measurements of the M13 procoat variants revealed an unexpected inverse relationship between Sec pathway load and fitness cost. Expression of the WT and 7N variants at full induction (Methods) imposed a significant fitness cost – reducing growth rate by over 30%. Unlike for other membrane proteins, the growth rate does not drop to zero, likely because the M13 procoat protein is much smaller than the average membrane protein (73 amino acids and one TMD versus 360 amino acids and five TMDs). In contrast, the 8N, 10N, and 11N variants showed no discernible growth impairment compared to the control without overexpression (Figure 4d) – the fitness cost essentially disappeared. This restoration of unrestricted growth with increased Sec pathway load is inconsistent with the hypothesis that Sec pathway saturation is the primary driver of the cost of membrane-protein overexpression. Since the overexpressed variants that fail to insert into the membrane will accumulate in the cytosol, this observation is also inconsistent with cytosolic aggregation of non-inserted membrane proteins causing the fitness cost. Rather, the data suggest that the reduced membrane insertion resulting from these manipulations reduces fitness costs (Figure 4c and d). Although the quantification of membrane insertion lacks the precision required to discern the subtle differences between the WT, 7N, and 8N variants (Methods), the effects of the 10N and 11N variants are strong (Figure 4c) and support the qualitative conclusion that fitness costs depend sensitively on protein insertion into the membrane. Therefore, these results are consistent with the hypothesis that the cost of membrane protein overexpression is primarily driven by severe spatial constraints within the membrane rather than Sec pathway overload.

### Overexpression and displacement of endogenous membrane proteins

To test if overexpressed membrane proteins displace endogenous ones, we developed a single-cell technique to track protein localization. We used a YFP-labeled endogenous membrane protein as a membrane localization reporter. When this endogenous membrane protein is displaced, we expect the YFP signal to shift from the membrane to the cytosol (Figure 5a). We chose AtpD-YFP and PstB-YFP as localization reporters because their membrane localization is particularly clear, facilitating the detection of changes (Methods). Although located in the cytosol, AtpD is the β-subunit of the membrane-localized ATP synthase complex (Boyer, 2002) and can be used as a reporter for the subcellular localization of this complex. PstB, a subunit of the high-affinity phosphate uptake system, was used as a second reporter because it is part of the membrane-embedded PstSCAB transporter complex (Xiao et al., 2026). We measured YFP signals in the different subcellular compartments using fluorescence microscopy and quantified the localization of the endogenous membrane protein using an *ad hoc* defined localization parameter (Figure 5a and b). While the values of this parameter varied from cell to cell and among proteins, they were predominantly positive for membrane proteins, whereas for a cytosolic control protein (TufB) the distribution was markedly shifted to negative values (Figure 5c). Taken together, this approach enables precise measurements of shifts in protein localization from membrane to cytosol.

**Figure 5:**
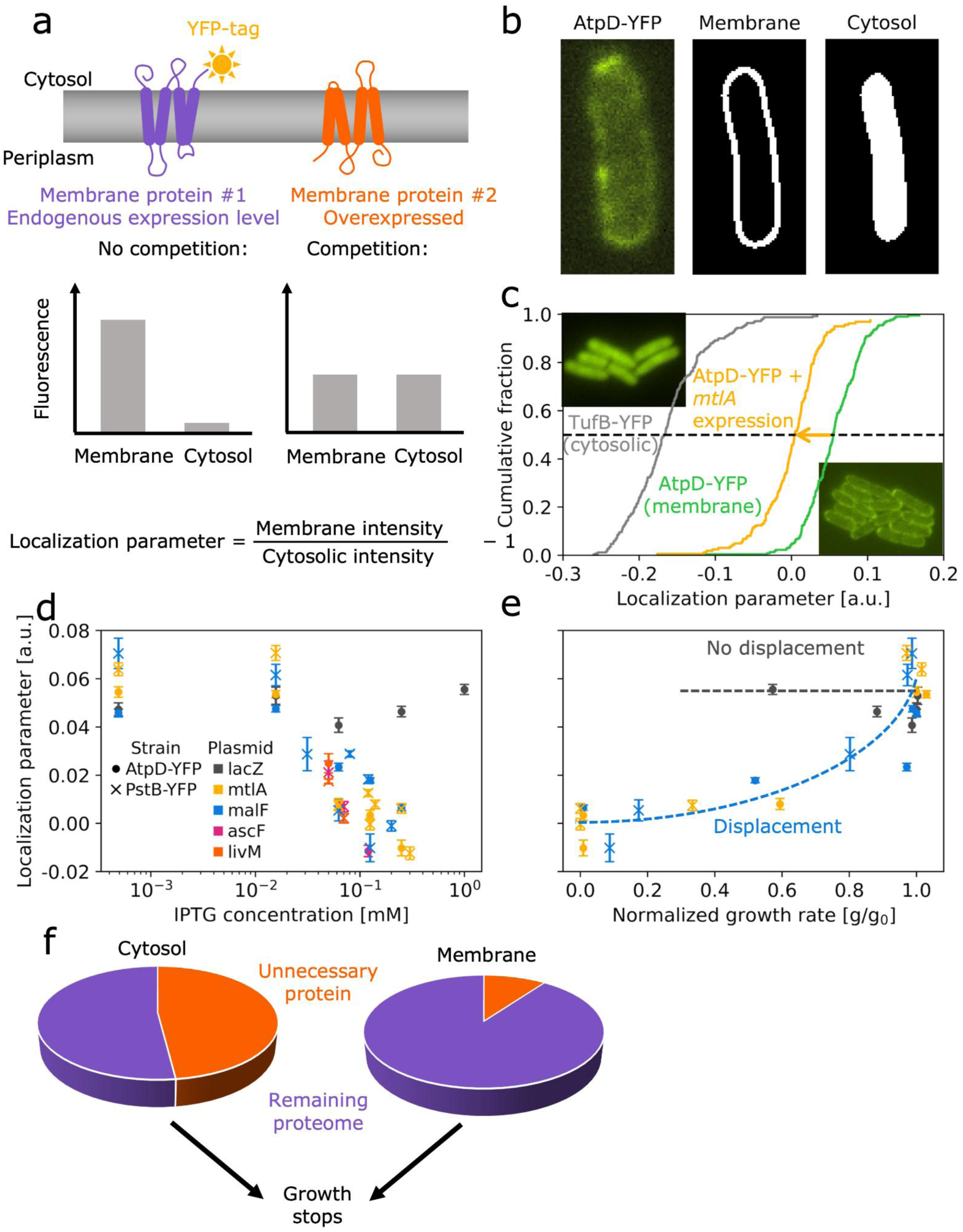
Overexpression of membrane proteins leads to the displacement of endogenous membrane proteins, which coincides with a collapse of the growth rate. **a)** Schematic of the assay used to detect protein displacement from the membrane. An endogenous membrane protein is fluorescently tagged, while a second membrane protein is overexpressed (Methods). If the proteins compete for membrane space, a fraction of the fluorescently tagged protein will remain in the cytosol instead of the membrane upon overexpression. **b)** Single-cell fluorescence images capture subcellular protein localization. Membrane and cytosolic regions are distinguished by image analysis (Methods). **c)** Cumulative distributions of localization parameters for different conditions and different proteins. TufB-YFP (gray) localizes to the cytosol (inset, top left) and has lower localization parameter values; AtpD-YFP (green) localizes to the membrane (inset, bottom right), and has higher values. Overexpression of MtlA (a membrane protein) in an AtpD-YFP strain (yellow) shifts the localization parameter distribution to lower values. **d)** Changes in the localization parameter of AtpD-YFP (circles) and PstB-YFP (crosses) upon overexpression of different proteins. Overexpression of membrane proteins (MtlA, yellow; MalF, blue; AscF, magenta; LivM, orange) decreases the localization parameter, reflecting the displacement of AtpD-YFP and PstB-YFP to the cytosol. Overexpression of a cytosolic protein (LacZ, gray) does not alter AtpD-YFP localization. Additional examples can be found in Supplementary Figure 11. **e)** Localization parameter from d) versus growth rate for MtlA and MalF overexpression. The decrease in growth rate coincides with a decrease in AtpD-YFP and PstB-YFP localization for strains overexpressing membrane proteins (MtlA, yellow; MalF, blue). Growth rates in this assay were precisely measured using bioluminescence (Methods). **f)** Pie charts of the estimated proteome mass fractions of overexpressed proteins at which growth stops for the cytosolic proteome (left) and the membrane proteome (right).

Overexpression of membrane proteins resulted in partial displacement of endogenous membrane proteins. We selected several membrane proteins covering different IC₅₀ values and with varying numbers of TMDs, including both even and odd numbers. We first overexpressed MtlA, a transporter of the sugar mannitol (Lee and Saier, 1983) with six TMDs, which was chosen as a representative membrane protein with a typical number of TMDs and because it is gratuitous under our growth conditions. Overexpressing MtlA consistently shifted the localization parameter to lower values (Figure 5c). The median membrane localization of the AtpD-YFP reporter was significantly reduced (*p* = 10^−35^, Mann–Whitney *U* test), but it still considerably exceeded that of a cytosolic control protein (Figure 5c). Similarly, overexpression of the membrane protein MalF, a subunit of the maltose transporter selected with the same rationale as MtlA, produced comparable reductions in AtpD-YFP membrane localization (Figure 5d). Similar results were obtained using PstB-YFP as a membrane-localization reporter (Figure 5d). This effect on membrane localization was further validated for overexpression of the membrane proteins AscF, LivM, AgaC, ArsB, PotC, and TrkH (Figure 5d and Supplementary Figure 11). In contrast, overexpression of the cytosolic protein LacZ as a control had no effect on membrane localization (Figure 5d). These results suggest that overexpression leads to a situation where the membrane capacity is exceeded and additional forced insertion of more membrane proteins leads to a partial displacement of the endogenous membrane proteome.

But does this displacement of endogenous membrane proteins cause the high fitness cost of membrane protein overexpression? To address this, we analyzed the relationship between membrane localization of the AtpD-YFP and PstB-YFP reporters and growth rate under conditions of steady-state exponential growth (Methods). The sharp decline in growth rate observed with increasing overexpression (Figure 3b) closely aligns with the loss of membrane localization, i.e., both occur at the same inducer concentration (Figure 5e), suggesting that the fitness cost begins to emerge when the displacement of endogenous membrane proteins becomes detectable. This pattern was consistent across different overexpressed membrane proteins and membrane-localization reporters, but did not appear for a cytosolic protein (Figure 5e). These findings support the hypothesis that the fitness cost of membrane protein overexpression arises from the partial displacement of the endogenous membrane proteome.

The sharp decline in growth rate upon membrane protein overexpression did not correlate with protein aggregation or inclusion-body formation in the cytosol, which may compromise the cell’s ability to maintain normal function (Bednarska et al., 2013). While we observed such protein aggregates via transmitted-light microscopy (Jong et al., 2017; Lindner et al., 2008) in case of cytosolic-protein overexpression, they were absent for membrane-protein overexpression (Supplementary Figure 12). Together with the data from the M13 procoat variants (Figure 4), these results largely rule out cytosolic protein aggregation as a major driver of the high fitness costs associated with membrane protein overexpression.

Our experimental data allow us to estimate the fraction of the membrane proteome that must be displaced to halt growth using two independent approaches. First, we converted the inducer concentration that causes growth arrest into a proteome fraction by comparing our LacZ data with published results (Supplementary Figure 7; Methods). Assuming a fixed capacity for proteins inside the membrane and no preferential selection for insertion of the overexpressed protein, our estimates suggest that growth arrest occurs when approximately 13% of the membrane proteome is displaced. Second, we used a computational simulation of the *E. coli* cell geometry as a spherocylinder to quantitatively interpret the experimentally observed localization parameters, taking into account the point-spread function in our microscopy setup. Consistent with our initial estimate, this analysis showed that displacing about 9% of the membrane proteome is sufficient to induce growth arrest (Supplementary Figure 13). The close agreement between these two estimates supports the notion that membrane protein capacity remains relatively constant. Notably, this threshold for growth arrest is much lower than the corresponding ∼50% displacement required for cytosolic protein overexpression (Scott et al., 2010) (Figure 5f), which is plausible because the membrane is highly crowded, has one less dimension, and much of the membrane proteome is essential.

### Dynamics of growth arrest and recovery

To further challenge the displacement hypothesis, we examined the dynamics of growth arrest upon membrane protein overexpression, as an increasing fraction of the membrane proteome is displaced. We followed single cells carrying the AtpD-YFP or PstB-YFP localization reporter in a microfluidic chamber using time-lapse microscopy (Methods). We first allowed bacteria to grow in normal growth medium for 1 hour before switching to a medium containing IPTG to induce overexpression of a membrane protein (MalF, MtlA, or LivM). For *malF* overexpression, the AtpD-YFP localization parameter decreased rapidly, following an approximately exponential relaxation on a time scale of 2.6±0.3 hours (Figure 6a). The growth rate followed a similar exponential decline on a time scale of 1.7±0.0 hours (Figure 6a); within about three generations, growth stopped entirely. Similar parallel declines in localization and growth rate occurred upon overexpression of *mtlA* (Figure 6b) and *livM* (Supplementary Figure 14a), as well as when PstB-YFP was used as localization reporter (Supplementary Figure 14c). Localization parameter and growth rate were highly correlated throughout these experiments (Figure 6c and Supplementary Figure 14b,d). This parallel decrease in localization parameter and growth rate is consistent with the view that the displacement of endogenous membrane proteins causes growth inhibition.

**Figure 6:**
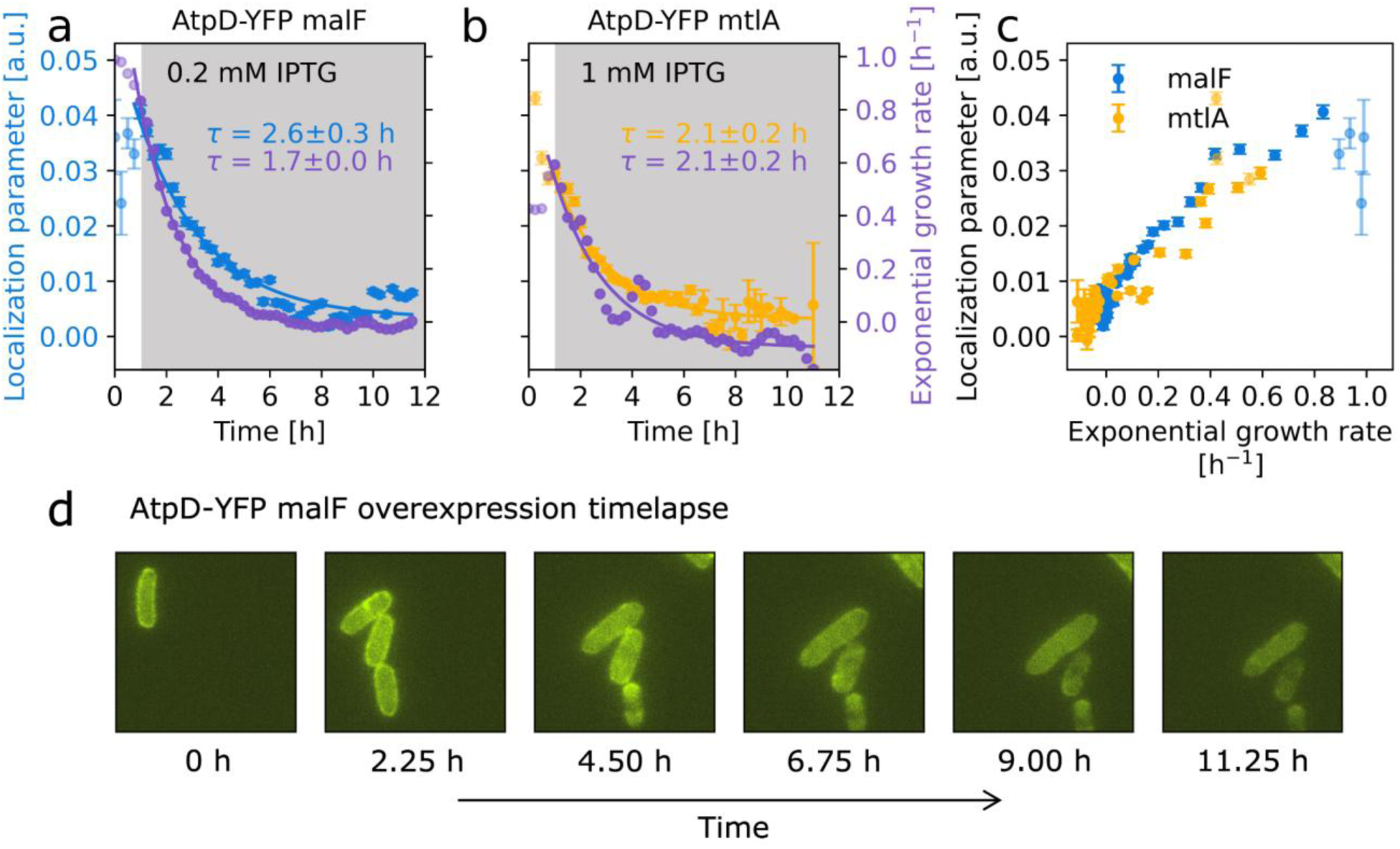
Displacement of membrane proteins occurs in parallel with the decrease in growth rate. **a)** Time course of the localization parameter for AtpD-YFP (blue) and the exponential growth rate (purple) upon MalF overexpression for single cells tracked in a microfluidic device (Methods). The gray-shaded area indicates the period during which IPTG (at 0.2 mM) was added to the medium. An exponential decay was fitted to the data (blue and purple lines) to determine the characteristic time scales, τ = 2.6±0.3 hours for the localization parameter and τ = 1.7±0.0 hours for the growth rate. **b)** As **a**, but for MtlA overexpression. Here, the characteristic time scale is τ = 2.1±0.2 hours for both the localization parameter and for the growth rate. **c)** Localization parameter for AtpD-YFP versus growth rate for different time points of the time-lapse experiments shown in **a** and **b**. Pearson correlation *r* = 0.92, *p* = 10^-19^ for MalF and *r* = 0.85, *p* = 10^-12^ for MtlA. Light blue and light yellow points represent early time points when the microfluidic chamber still contains relatively few cells. **d)** Time-lapse fluorescence images of single cells from the experiment in **a**. Supplementary Figure 14 provides additional examples of this effect for other membrane proteins.

Partial membrane proteome displacement trapped most bacteria in a non-growing state. To test how bacteria recover from overexpression, we switched the microfluidic device back to inducer-free growth medium after 5 hours of *malF* overexpression. Remarkably, the vast majority of bacteria (148 out of 162) failed to resume growth even after the inducer was removed (Supplementary Figure 15). A small fraction did recover and began to crowd the microfluidic chamber after about 10 hours, preventing the observation of non-growing cells for longer periods of time. For comparison, typical lag times for *E. coli* cells recovering from stationary phase are about 1 hour (Fridman et al., 2014). Thus, the partial displacement of endogenous membrane proteins by overexpression, which occurs in as little as 5 hours (Figure 6a and b; Supplementary Figure 14a,c), causes a persistent perturbation of the membrane proteome, trapping cells in a nearly irreversible state of growth arrest. This observation further supports the displacement hypothesis because, once inserted, the space occupied by a membrane protein is irreversibly lost to the cell; in contrast, fitness costs due to translocation pathway saturation should be transient and rapidly decrease once membrane-protein overexpression ceases.

## Discussion

Our study provides a quantitative genome-wide analysis of the fitness costs of protein overexpression (Figure 1), revealing extremely high costs (Figure 2) and sensitivity to expression level for membrane proteins (Figure 3). Using engineered membrane protein variants, we have shown that these costs are not due to saturation of the membrane translocation machinery (Figure 4). Instead, the costs primarily arise from the limited membrane space being overwhelmed by the overexpressed protein. The fitness cost increased with the amount of protein inserted (Figure 4), with protein hydrophobicity (Supplementary Figure 8), and with the number of TMDs (Figure 2d), a close proxy for the membrane space occupied. We provided evidence that this fitness cost is mainly due to the displacement of endogenous membrane proteins from the crowded membrane (Figure 5). Single-cell experiments revealed that this displacement occurs in tandem with a sharp decline in growth rate (Figures 5 and 6). Our analysis indicates that displacement of as little as 10% of the endogenous membrane proteome is sufficient to halt growth (Figure 5f), leaving the bacteria behind in a perturbed state from which they can hardly recover.

It is plausible that this growth arrest is essentially irreversible because bacteria lack effective mechanisms to degrade or otherwise remove excess proteins from the membrane. Although some integral membrane proteases exist, such as HtpX and FtsH (Sakoh et al., 2005), they are either downregulated upon membrane protein overexpression (HtpX) (Wagner et al., 2007) or target only specific proteins (FtsH) (Ito and Akiyama, 2005). Therefore, once the membrane is overloaded with useless “junk” proteins, non-growing bacteria struggle to resume growth, even after the overexpression is stopped.

It has often been assumed that the main problem with membrane protein overexpression is the saturation of the membrane translocation machinery. This assumption has inspired efforts to boost membrane protein yields by overexpressing components of the Sec pathway, but these have been largely unsuccessful. One reason for this is that most Sec components are themselves membrane localized, which imposes a high fitness cost (Steinberg et al., 2018). Our results imply that this approach has limited promise: our data consistently show that the limiting factor for membrane protein expression is not the Sec pathway, but the membrane’s physical capacity. Our data suggest that the cost of a TMD passing through the membrane, possibly due to the required energy, provides only a small additional contribution to the fitness cost of membrane protein expression, as reflected by slightly higher costs for proteins with an odd number of TMDs (Figure 2d,e). Therefore, a more promising strategy to increase yields is to increase membrane capacity by modifying the physical properties of the membrane. This could be achieved by increasing lipid production (Mercier et al., 2013) or by manipulating other membrane properties such as its lipid composition. Consistent with this conclusion, recent work in eukaryotes has shown that enhancing membrane fluidity can increase membrane protein capacity and ultimately improve cellular fitness under overexpression conditions (Kanonenberg et al., 2019; Pichler and Emmerstorfer-Augustin, 2018). Another approach could be to reduce the expression of non-essential genes that occupy a large membrane area to free up membrane space for the overexpressed protein. Developing efficient ways to increase membrane capacity for the overexpressed protein is a promising direction for advancing this field.

In addition to its practical relevance in increasing membrane protein yields, the displacement hypothesis makes several predictions that are relevant to basic research. One interesting direction is altering membrane fluidity. While this could be achieved by changing the temperature or using chemicals (Hodzic et al., 2012), these perturbations typically impact numerous cellular processes, introducing confounding factors that are challenging to discern. Another possibility is manipulating the expression of integral membrane proteases, which could alter protein density in the membrane. This could free up or limit membrane space further, thereby altering the extent of overexpression of other membrane proteins required to cause growth collapse due to exhaustion of membrane capacity. However, characterized integral membrane proteases often degrade only specific membrane proteins or membrane proteins in abnormal states (Ito and Akiyama, 2005), so their overall effect on an overcrowded membrane may be limited. Properly conducting the necessary experiments is challenging and beyond the scope of this study, but future work could test these predictions to further corroborate the displacement hypothesis.

Our findings also have implications for the evolution of antibiotic resistance, especially for mechanisms involving efflux pumps. Resistance often arises from spontaneous mutations that increase the expression of large membrane protein complexes, such as the AcrAB-TolC multidrug efflux pump in *E. coli* (Toprak et al., 2012). Such overexpression comes at a high cost: as the membrane capacity is approached, an abrupt growth collapse occurs (Figure 3b). This imposes a hard ceiling on resistance evolution via efflux pump overexpression: at some point, the fitness costs of this mechanism become unsustainable and outweigh the benefits, forcing evolution to seek alternative ways to increase resistance. Notably, this may explain the observed pattern of spontaneous resistance evolution to tetracyclines in *E. coli*, which often plateaus after modest increases mediated by increased expression of the AcrAB-TolC efflux pump (Toprak et al., 2012), presumably due to the lack of readily accessible alternative resistance mechanisms. In contrast, resistance to chloramphenicol readily evolves to higher levels via increased expression of the MdfA efflux pump (Toprak et al., 2012), a smaller and thus less costly membrane protein. Note that we cannot directly compare the expression costs of MdfA and AcrAB-TolC because our quantitative overexpression dataset only includes the individual components (AcrA, AcrB, and TolC), rather than the entire AcrAB-TolC multiprotein complex. A promising direction for future studies is to extend our systematic investigation of overexpression costs to include such multiprotein complexes. It will be interesting to explore the extent to which the expression costs of different resistance proteins can help to predict preferred mutational trajectories in resistance evolution.

While our study highlights the extreme cost of membrane-protein expression, the quantitative genome-wide fitness cost data we generated is a powerful resource for exploring many other aspects of bacterial physiology. Notably, transcriptional regulators and other DNA-binding proteins exhibit some of the highest expression costs in the genome (Figure 2). It is certainly plausible that perturbations in transcriptional regulation strongly affect fitness. One possible explanation is that DNA-binding proteins may face similar constraints as membrane proteins, where a limited cellular resource—in this case, binding space on the DNA —becomes a bottleneck. However, uncovering the mechanisms behind this phenomenon will require a dedicated study focused on this topic. We hope that our work will inspire similar quantitative studies on protein overexpression in other organisms. Such quantitative information on expression costs, combined with the wealth of information already available on proteins (Karp et al., 2023; The UniProt Consortium, 2025), has the potential to advance both basic and applied research in microbiology and beyond.

## Materials and Methods

### Experimental methods

#### Bacterial strains

The genome-wide measurement of overexpression costs (Figures 1–3) was performed using strains from the ASKA overexpression library (Kitagawa et al., 2006). The library was transferred from 56 96-well plates to fourteen 384-well plates, with 14 to 145 transcription-only controls added to each plate. For these controls, the plasmid pAA031 was used, in which the open reading frame was cleanly deleted (Kavčič et al., 2020a).

M13 procoat overexpression strains (Figure 4) were constructed by transforming *E. coli* K-12 BW25113 with the pCA24N (ASKA) plasmid containing the open reading frame of the respective M13 procoat variant. To prepare these plasmids, the original empty ASKA plasmid (Kitagawa et al., 2006) – with GFP excised from the reading frame – was digested with StuI, phosphorylated, and then ligated with the M13 procoat sequence (as described in (Kitagawa et al., 2006)). The open reading frames of the WT and mutant M13 procoat variants (7N, 8N, 10N, 11N) were generated by commercial gene synthesis (Eurofins Genomics) and cloned into the pEX-A128 vector backbone. They were then subcloned into pCA24N, as described above. All constructs were verified by Sanger sequencing.

For the construction of the AtpD-YFP strains (Figure 5 and 6), we started with the DY330 AtpD::YFP::Cam^R^ strain from (Taniguchi et al., 2010). In this strain, the Cam^R^ resistance cassette was replaced by the Amp^R^ resistance cassette by homologous recombination (Lynn C. Thomason, James A. Sawitzke, Xuan Li, Nina Costantino, Donald L. Court, 2005). The modified AtpD::YFP::Amp^R^ sequence was then transferred to the wildtype BW25113 strain (Baba et al., 2006) by P1 transduction (Lennox, E.S., 1955). The resulting BW25113 AtpD::YFP::Amp^R^ strain was then transformed with overexpression plasmids containing open reading frames for membrane proteins (*malF* and *mtlA*) and the cytosolic control protein (*lacZ*). *malF* and *mtlA* were selected because they are neutral, unnecessary membrane proteins encoding transporters for carbon sources absent in Lysogeny Broth (LB) medium, and because they have a typical number of transmembrane domains (8 and 6, respectively). In addition, these strains were transformed with the pCS-λ plasmid, containing the bacterial *luxCDABE* operon driven by the constitutive λ-PR promoter to facilitate precise bioluminescence-based growth-rate measurements (Kavčič et al., 2020b).

#### Growth media

All colony growth assays (Figures 1–3, Supplementary Figure 1) were performed on omnitrays (VWR, 734-0490) filled with 43 mL solid LB agar medium (Sigma-Aldrich, L2897). Expression levels were measured in minimal M9 medium (1 × M9 salts (Sigma-Aldrich, M6030), 2 mM MgSO_4 (_Sigma-Aldrich, M7506), 0.1 mM CaCl_2 (_VWR, 22317), supplemented with 4 g/L glucose (Sigma-Aldrich, G8270) and 1 g/L amicase (Sigma-Aldrich, A2427)). In addition, Triton X-100 (Sigma-Aldrich, T8787) was added at 0.001% (v/v) to reduce surface tension in the microplate wells; this had no detectable effect on growth or gene expression. For all other experiments (Figures 4–6, Supplementary Figures 3, 7, 12 and 15) liquid LB Lennox medium (Sigma Aldrich, L3022) was used.

The antibiotics used for selection were chloramphenicol (Sigma-Aldrich, C0378), ampicillin (Sigma-Aldrich, A9518) and kanamycin (Sigma-Aldrich, K4000). The chloramphenicol stock solution was prepared by dissolving it in ethanol, while the ampicillin and kanamycin stocks were dissolved in pure water. All stock solutions were filter-sterilized and stored at −20°C in the dark. Aliquots were thawed at room temperature before use. When necessary, selection was performed at 50 μg/mL for kanamycin and at 100 μg/mL for ampicillin. Selection for overexpression plasmids was performed at 35 μg/mL of chloramphenicol. A filter-sterilized solution of 1 M IPTG (VWR, 437144N) in water was used as the stock solution. IPTG was stored at −20°C in the dark and aliquots were thawed at room temperature before use. For all strains carrying overexpression plasmids, chloramphenicol at 35 µg/mL was also used during the experiments to ensure maintenance of the plasmid, while other selection markers were omitted.

#### Colony growth

LB agar plates were prepared in batches for each of the twelve IPTG concentrations and stored at 4°C for six to thirty days. IPTG was added to liquid agar after cooling to 60–65°C, consistent with standard protocols. Because IPTG is heat-sensitive and is not typically autoclaved, slight degradation during this step cannot be excluded. Prior to the experiment, the plates were equilibrated to room temperature for 2 hours. In each individual run of the experiment, one of the fourteen 384-well ASKA overexpression plates was measured across twelve IPTG concentrations. The 384-well plate was thawed at room temperature for one hour, after which the strains were inoculated onto the agar plates in quadruplicate using a colony-processing robot (BM3-BC, S&P Robotics). A built-in camera (Canon, EOS 700D) captured time-lapse images of the plates every 15 minutes for 26 hours. Inoculation and imaging were performed in a room with controlled temperature and humidity (30°C, 50%). All images were stored in CR2 raw format.

#### Colony biomass

To measure colony biomass over time and compare it to colony intensity (Supplementary Figure 1), we inoculated ten LB agar plates with six colonies each using the colony processing robot. The plates were incubated at 30°C and 50% humidity, and biomass and intensity were measured for three colonies every 30 to 40 minutes for 9 hours. For the colony intensity measurement, the plates were imaged using the above mentioned built-in camera. Images were analyzed as described below. For the biomass measurement, each colony was carefully cut from the agar using a small (ca. 1 cm²) cookie cutter and transferred to a Falcon tube filled with 15 mL isotonic solution (Isotone II Diluent, Beckman Coulter, 8448011). The tube was vortexed for 30 seconds to detach the cells from the agar pad while leaving the pad itself intact. The solution was then filtered (Sigma-Aldrich, SLAP02550) to remove any small agar fragments, and the total cell volume and cell count were then determined using a Coulter counter (Multisizer 3, Beckman Coulter). The biomass was determined by multiplying the measured cell volumes with a mass density of 0.34 g/mL (Ridgway et al., 2008).

#### Growth rate of M13 procoat mutants

To measure the fitness cost of the M13 procoat variants (Figure 4), an IPTG gradient with eight concentrations (ranging from 0.008 mM to 1 mM in logarithmic steps) was prepared along the columns of a clear 96-well microtiter plate (VWR, 732-2719). Bacteria were inoculated from fresh overnight cultures (ONC) at a 1:1,000 dilution, and each strain and condition was tested in quadruplicate. Plates were incubated in a plate reader (Biotek, Synergy Neo2) at 30°C and 807 rpm, and absorbance at 600 nm (A_600)_ was measured every 10 minutes for 20 hours. Growth rates were calculated from the resulting growth curves as described below.

#### Expression-level measurements

We performed two related expression-level measurements using ASKA strains with C-terminal GFP fusions. For Supplementary Figure 4, we selected 41 strains spanning the IC₅₀ range of the genome-wide overexpression dataset, including both low-cost and high-cost proteins with different subcellular locations. This experiment was designed to test whether IPTG concentration produces a broadly consistent change in expression level across proteins with different overexpression costs. Each strain was measured in two independent replicate experiments. The data from this experiment, including strain identities, are available in Supplementary Table 2.

For Supplementary Figure 5, we performed a separate experiment using 78 GFP-tagged ASKA strains to compare growth inhibition as a function of directly measured GFP fluorescence for membrane and cytosolic proteins. These data were used to estimate E75, defined as the expression level at which growth decreased to 75% of the growth rate without inducer. This experiment was designed as an additional validation that membrane-protein overexpression causes growth inhibition at lower measured expression levels than cytosolic-protein overexpression. Unlike the measurements shown in Supplementary Figure 4, these measurements were not performed in duplicate. The data from this experiment, including strain identities, are available in Supplementary Table 3.

For both experiments, an IPTG concentration gradient was established in minimal M9 medium across twelve 96-well microtiter plates, using the same concentrations as in the colony growth assay and 200 µL of medium per well. The strains were inoculated into these plates at a 1:1,000 dilution. Plates were incubated in a shaking incubator (Liconic, STX44) at 30°C and 97% humidity to minimize evaporation. A_600 a_nd GFP fluorescence were measured every 20–30 minutes using a plate reader (Biotek, Synergy H1) integrated into a robotic system (HighRes biosolutions MC642). Fluorescence measurements were performed with excitation at 485 nm and emission at 528 nm (both BW 20nm, Biotek).

Expression levels were determined as previously described (Angermayr et al., 2022; Mitosch et al., 2017). Briefly, each plate contained a control strain with the pAA031 control plasmid lacking GFP expression, grown in parallel with GFP-expressing strains. For all strains, the exponential growth phase was identified and the GFP signal of the control strain was subtracted from the GFP signal of the GFP-expressing strains at the corresponding A_600._ Linear interpolation (*numpy.interp*, NumPy 1.21.6) was used to interpolate GFP values between the two closest A_600 m_easurements. Expression level was calculated as the average GFP/A_600 r_atio during exponential growth, for A_600 v_alues between 0.03 and 0.15.

#### Membrane localization

To quantify AtpD-YFP localization at different overexpression levels of *malF*, *mtlA*, or *lacZ*, microscopy images were captured during the exponential growth phase (Figure 5). We prepared a 12-concentration IPTG gradient by two-fold serial dilution, ranging from 1 µM to 1 mM in eleven steps and one batch without IPTG. This concentration gradient was added to the columns of two 96-well microtiter plates, in which bacteria from fresh ONCs were inoculated at different dilutions. A white 96-well plate (VWR, 732-2697) was prepared with bacteria at a 1:10,000 dilution for bioluminescence-based growth measurements, with measurements taken every 5 minutes for 24 hours using a plate reader (Biotek, Synergy Neo2). A second clear 96-well plate was inoculated with bacteria at a dilution of 1:500 to 1:1,000 and A_600 m_easurements of this plate were taken every 10 minutes for 8 hours in a separate plate reader (Biotek, Synergy H1).

A subset of samples from the second plate was selected for microscopy based on IPTG concentration and the observed growth inhibition. When the A_600 r_eached a value between 0.05 and 0.1, the samples were prepared for imaging. A 1.5 µL drop of the sample was transferred to an agar pad and the agar pad was placed on a coverslip such that the bacteria were “sandwiched” between the coverslip and the agar pad. Microscopy images were captured using a 100× oil objective with an EMCCD camera (Hamamatsu) on a Nikon Eclipse Ti-E inverted microscope with a Lumencor LED light engine, controlled by NIS Elements software. Phase contrast and YFP fluorescence images were acquired. Excitation wavelengths for YFP were CWL/FWHM 513/17 nm and emission wavelengths were dichroic LP 520 nm, CWL/BW 542/27 nm. In addition, a Z-stack of eight steps at 0.15 µm intervals was performed in the YFP channel to ensure accurate focal plane imaging for the localization measurements.

#### Microfluidics and time-lapse microscopy

For time-lapse microscopy experiments, we used a microfluidic device (CellASIC ONIX, Merck Millipore), which allows bacterial growth in microcolonies and enables rapid switching between different inlets, allowing complete media exchange within minutes (Mitosch et al., 2017). Bacteria were inoculated from frozen glycerol stocks at a dilution of 1:1,000 to 1:2,000 and grown to an A_600 o_f 0.05 to 0.1. Cultures were then diluted 1:100 and loaded into a microfluidic chamber preheated to 30°C, typically resulting in spatially separated single cells within the chamber. All experiments were performed in an incubator box at 30°C. Data acquisition began 30 minutes after loading and images were acquired every 15 minutes for 11 hours using the same equipment as described in the previous section. At each time point, a phase-contrast image and a Z-stack (5 images at 0.15 µm intervals) in the YFP channel were acquired. IPTG induction was started one hour after loading. In IPTG removal experiments, the medium was switched back to medium without IPTG 5 hours after induction.

### Computational Analysis

#### Colony growth rate

##### Image segmentation

The images obtained by the colony-processing robot were read using the RawPy Python package (version 0.15.0) and were cropped to include only the area containing the colonies. They were then automatically segmented into tiles, each containing a single colony. This segmentation was achieved by calculating intensity profiles along both axes, smoothing these profiles, and identifying intensity peaks as local maxima. The data were smoothed by convolving them with a normalized Blackman window of a specified width. The smoothing width was iteratively increased until the number of detected local maxima matched the expected layout. These local maxima defined each colony’s center coordinates, and the final smoothing width typically ranged from 20 to 30 pixels. Segment boundaries were defined as the midpoints between adjacent colony-center coordinates.

##### Speckle detection

Speckles in the background image were detected using three complementary methods: identification of background pixels with high intensity gradients, unusually high intensities, or sudden extreme temporal intensity changes. This is achieved by thresholding, with suitable thresholds determined from previous comparable experimental data. The detected regions were dilated by a disk of radius 3 pixels and excluded from all further calculations. Note that subsequent data analysis and conclusions are largely unaffected by the details of speckle detection.

##### Colony area and background area in each segment

To determine the colony area, a circular area with a radius of ten pixels around the colony centers was treated as the inner part of the colony. The radius of this circle was selected to match the pin size of the robotic pin tool used for colony inoculation. For the background area, the colony area was dilated by a disk of radius 8 pixels (resulting in an 18-pixel-radius circle around the colony center); the remainder of the segment was used as the background area.

##### Colony intensity curves

For each timepoint and segment, the mean pixel intensity within the colony and background areas, respectively, was calculated and the background intensity was subtracted from the colony intensity. Before calculating growth rates from these curves, each curve was again normalized to the mean of the first eight data points (corresponding to 2 hours). Similar approaches for generating growth curves from colonies on solid agar have been presented previously (Kamrad et al., 2020; Kritikos et al., 2017; Levin-Reisman et al., 2010; Takeuchi et al., 2014; Zackrisson et al., 2016).

#### Growth rate calculation

To calculate the exponential growth rate of microbial growth curves, we selected the data within defined time and absorbance/bioluminescence/intensity windows. The specific windows used were time: between 2 and 15 hours; A_600:_ between 0.005 and 0.1; bioluminescence: between 10^2^ and 10^5^; intensity: between 10^2^ and 10^3^. We applied a sliding linear fit over intervals of ten data points to estimate exponential growth rates from the slope of the logarithmic values. The maximum value over all intervals was taken as the exponential growth rate. If the lower threshold for the values (e.g., 0.005 for A_600)_ was not exceeded, the growth rate was set to zero. Growth rates showed modest variation between batches, with virtually no edge or neighbor effects observed on plates (Supplementary Figure 2).

#### Overexpression dose-response curves

For each intensity growth curve, we also determined the time point of maximum growth. This was calculated by using the same algorithm as for the growth rate, but with an output of the midpoint of the fitted time interval instead of the rate. If the measured growth rate of the sample was within 80% of the wildtype growth rate and this time point was later than 15 hours, the replicate was excluded from the analysis, as such late growth is most likely due to contamination or escape mutants. Similarly, if the growth rate increased by more than 30% of the wildtype growth rate with increasing inducer concentration, the increased growth rate was set to the minimum value measured in the dose-response curve because such abrupt increases in growth rate with overexpression are implausible and likely due to contamination or escape mutants.

If one of the four replicates deviated strongly from the other three, it was excluded from the analysis. The criterion for detecting such outliers was that the variance of the growth rates variance of the replicates was greater than 0.01 h^−2^; the outlier was detected as the replicate furthest from the mean. For each strain, all growth rates along the IPTG gradient that were not excluded by the above criteria were normalized to the mean growth rate of the lowest three IPTG concentrations (including zero), which generally had no detectable effect on growth, and fitted to a Hill function *y*(*x*) = 1/[1 + (*x*/IC_50)_*^n^*] using SciPy’s (1.7.3) `optimize.curve_fit`. The IC_50 a_nd Hill coefficient *n*, which were used for further analysis of the data, were obtained from this fit. In the experiments, we used inducer concentrations up to 1 mM and therefore set the maximum IC_50 v_alue for the Hill fit to 2.5 mM. This ensured that even dose-response curves showing minimal or no inhibition were assigned a consistent IC_50 v_alue, enabling rank-based statistical analysis across the entire gene set.

#### Gene-ontology enrichment

The `enrichr` function of GSEApy 1.1.3 was used for the gene-ontology enrichment analysis. The combined score, that we use as a measure for enrichment in Figure 2a and Supplementary Figures 6 and 9, is calculated as the product of the odds ratio and the negative logarithm of the p-value, Combined score = odd′s ratio × −log(p).The necessary data for the GO terms of each gene was collected from the UniProt database using the information on *E. coli* K12 (The UniProt Consortium, 2025). The necessary structure on the gene ontology tree was retrieved through the go-basic.obo file from the GO knowledgebase (Ashburner et al., 2000; The Gene Ontology Consortium et al., 2023).

#### Protein features and classifications

All protein features (such as subcellular localization, length, number of transmembrane domains, etc.) were obtained from the UniProt database (The UniProt Consortium, 2025). This protein information was matched to the ASKA library using the ordered locus name or the JW identifier.

Hydrophobicity values for individual amino acids were obtained as normalized sidechain hydrophobicity from (Monera et al., 1995).

The alternating local IC_50 t_rend based on the parity of the number of TMDs was validated using a bootstrapping approach (Beran, 2008). To do this, we resampled the IC_50 d_istributions for each TMD group, calculated the medians of the resampled data, and evaluated whether an alternating pattern was present compared to the running average of the new data. To quantify this pattern, we multiplied the local IC_50 t_rends by the alternating function (-1)^m^, where m represents the number of TMDs, and summed the resulting values. A positive sum indicates an alternating trend consistent with the proposed direction (higher local IC_50 v_alues for even numbers of TMDs and lower for odd numbers), while a negative sum indicates the opposite. This resampling procedure was repeated 10,000 times, and the same parity trend was observed in 96% of all iterations.

#### Microscopy image analysis

Cell contours were identified using phase-contrast images. To reduce noise, the images were first smoothed by convolution with a normalized Blackman window of 8 pixels width. Next, a global threshold was determined from the smoothed image using a minimum-threshold algorithm (`filters.threshold_minimum`, skimage, 0.15.0) to distinguish cells from background. Based on this threshold, a binary mask was created and then dilated by a disk of radius 2 pixels to improve cell-boundary detection. Contours in the binary mask were identified using an edge-tracing algorithm (`measure.find_contours`, skimage, 0.15.0), producing a set of candidate cell boundaries. Each identified contour was then evaluated based on its size and shape characteristics: Contours with a minimum contour length of 100 pixels (minimum cell size) and an area-to-contour-length ratio between 2 and 10 px^2^/px were retained; contours near the image borders or outside the defined ranges were excluded. For each cell, the membrane area was defined as the region within the cell contour extending five pixels inward, while the remaining inner cell area was defined as the cytosolic area (Figure 5b). Similar image-analysis methods to this have been presented previously (Goudsmits et al., 2016).

The width of the membrane area was iteratively determined for a few samples of the AtpD-YFP strain without overexpression to maximize the localization parameter. The localization parameter for individual cells was calculated as the ratio of the mean YFP intensity in the membrane area to the mean YFP intensity in the cytosolic area, subtracted by one. From the Z-series of images acquired in the YFP channel, the Z-plane with the highest localization parameter was selected for each cell, as localization decreased outside the focal plane. To compare changes in membrane localization across different conditions and for different overexpressed proteins, the median of the observed localization-parameter distributions was used, as it is more robust against extreme values that can sometimes occur due to failure of the automatic image analysis.

#### Inserted protein fraction of M13 procoat variants

To estimate the membrane-inserted fraction of the M13 procoat variants, we performed densitometric analysis of published gel images from (Hariharan et al., 2021). The original image files were not available to us, therefore the analysis was performed on the images as published in the article. The WT, 7N, 8N, and 10N variants were quantified from Figure 1 of (Hariharan et al., 2021), and the 11N variant was quantified from Figure 2 of the same study.

In these experiments, the cytoplasmic region of M13 procoat was extended by 103 amino acids of SP1 (also known as Lep) to allow immunoprecipitation with Lep antiserum. Three distinct bands were observed on the protein gels of the M13 procoat mutant proteins in Figure 1 and 2 of (Hariharan et al., 2021): The upper band corresponds to non-translocated protein retained in the cytosol, whereas the lower processed bands correspond to membrane-translocated protein after signal peptidase cleavage and, where applicable, additional proteinase K cleavage. We focused on conditions in which both YidC and SecE were present, matching the physiological conditions in our experimental strains.

For each gel image, lane boundaries were determined from smoothed horizontal intensity profiles and used to crop individual lanes. Vertical intensity profiles were then generated for each lane. Because of the limited resolution and noise of the published images, band positions were identified manually rather than by automated peak detection. For each lane, the signal corresponding to inserted protein was divided by the total signal detected, corresponding to inserted plus non-inserted protein. When both signal-peptidase and proteinase-K-treated conditions were available, the estimated inserted fractions were averaged.

This analysis should be interpreted as an approximate quantification of the published data, not as precise densitometry of raw gel images. Published images may be affected by reduced resolution, compression, contrast adjustment, and other image-processing steps. Therefore, while the resulting values are sufficient to capture the qualitative reduction in membrane insertion across the variant series, they do not allow for the robust discrimination of small differences between closely spaced variants, especially WT, 7N, and 8N.

#### Localization parameter from computational simulation of cell geometry

A three-dimensional spherocylindrical rod model was created to simulate membrane and cytosolic protein distributions within a bacterial cell and the resulting fluorescence microscopy images. The rod was generated by positioning a sequence of spheres along a central axis, with the sphere radius and central axis length chosen to match typical cell dimensions. Specifically, each pixel represented 12.5 nm and we used a rod diameter of 40 pixels (500 nm) and a length of 90 pixels (1.125 µm). To define the cytosolic region, the rod radius was reduced by 3 pixels to account for the membrane and periplasmic space (Supplementary Figure 13c), assuming 7.5 nm for the width of the outer and plasma membranes (Milo et al., 2010)(BNID 109060), respectively, and 17.5 nm for the periplasmic space (Milo et al., 2010)(BIND114122). The membrane was defined as the outer layer of the rod with a width of 1 pixel (Supplementary Figure 13a).

To simulate the optical properties of the fluorescence microscope, the *z*-sum of the protein distribution was smoothed by convolution with a Gaussian distribution with a standard deviation σ_xy o_f 7.6 pixels to mimic the point-spread function (PSF) for our imaging setup (Supplementary Figure 13b and d). The value of σ_xy w_as calculated using the Rayleigh criterion with a YFP emission wavelength of 530 nm and a numerical aperture of 1.45, as used in our experiments. Next, we defined masks for the membrane and cytosolic regions to evaluate the localization parameter of the simulated images, analogous to the experimentally acquired images. A cross-sectional slice of the rod at its midpoint along the *z*-axis was used to define regions corresponding to the membrane and cytosol. Mask boundaries and widths were iteratively tested to maximize the localization parameter, defined as the mean fluorescence intensity ratio between the membrane and cytosolic regions, subtracted by one, just as for the experimental images. The cytosolic mask was defined as the area within 5 pixels from the longitudinal axis of the simulated cell and the membrane mask as the area between 11 and 14 pixels of distance from the longitudinal axis. After identifying these mask parameters, the final membrane and cytosolic masks were applied to the simulated microscope images. Different protein distributions in the membrane and cytosol were simulated, keeping the total protein amount constant and the distributions within the two compartments homogeneous, while varying the membrane-to-cytosol signal ratio from 10^5^ to 10^-5^ (Supplementary Figure 13e). These distributions were convolved with the Gaussian PSF and summed over the *z*-axis to generate the final simulated microscope image. The localization parameter was then calculated for each simulated image using the optimized membrane and cytosol masks (Supplementary Figure 13e).

## Acknowledgments

We thank Adam Palmer and Yuval Mulla for feedback on the manuscript; Adriana Espinosa-Cantu and Booshini Fernando for help with cloning; and Karin Schnetz, Joachim Krug, and the entire Bollenbach group for fruitful discussions. This work was supported in part by German Research Foundation (DFG) standalone grant BO 3502/2-1 and Collaborative Research Centre (SFB) 1310. SAA was supported by a Marie Skłodowska-Curie Individual Fellowship, grant agreement No. 707352, from the European Commission.

## Author contributions

Conceptualisation: J.M., S.A.A. and T.B.; Investigation: J.M. and N.G.; Statistical Analysis: J.M. and G.A., Computational modeling: J.M.; Writing original draft: J.M.; writing - review and editing: J.M., S.A.A., G.A., T.B.; Supervision: T.B.; Funding acquisition: T.B.

## Supplementary Figures

**Supplementary Figure 1:**
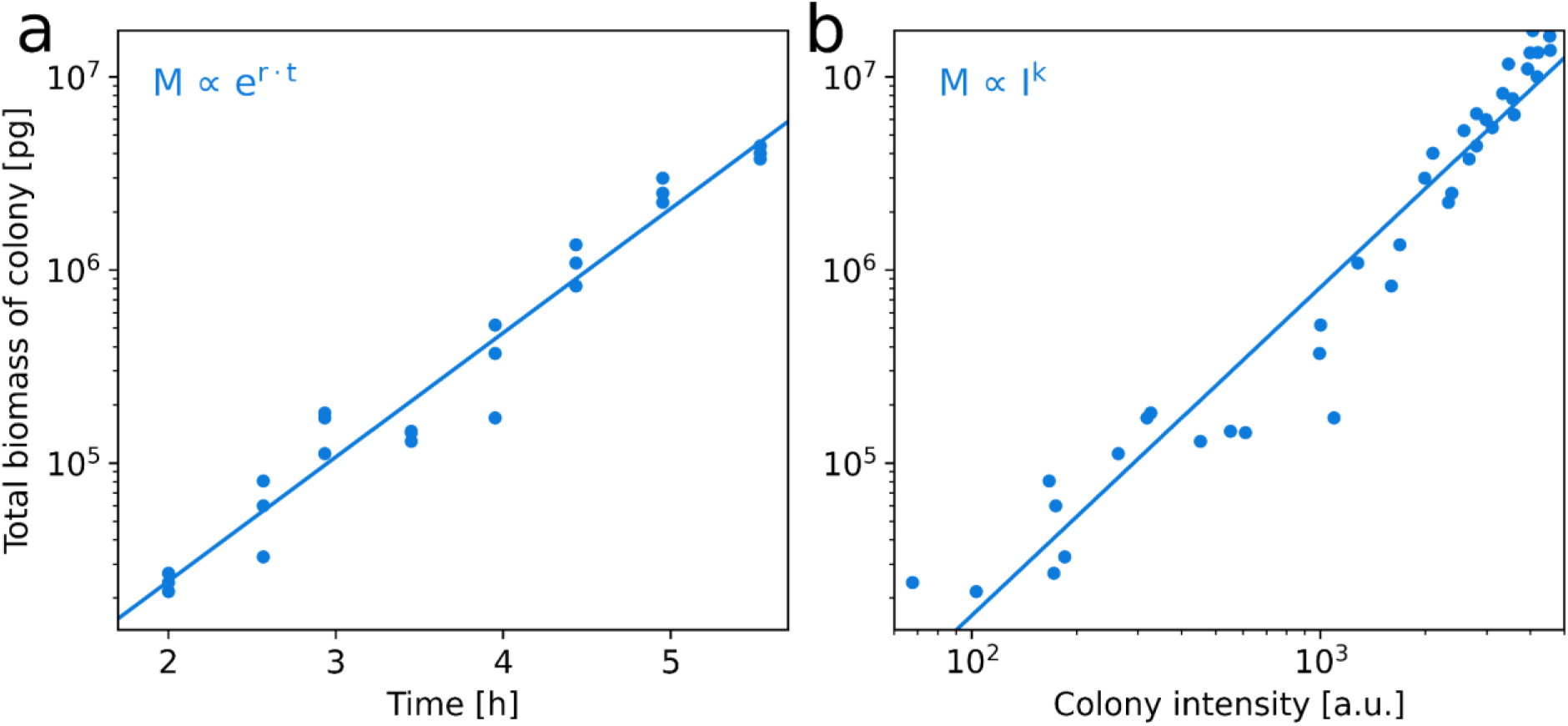
Colony intensity is a good proxy for colony biomass, which initially grows exponentially. **a)** Total biomass of *E. coli* colonies as a function of time. Replicate colonies were inoculated at the same time. Each data point represents a single colony for which biomass was determined at the respective time. For this we measured the total cell volume and calculated the total biomass assuming a constant dry mass density (Methods). **b)** Colony biomass vs. intensity at different time points. During the exponential growth phase (between 2×10^4^ pg and 5×10^6^ pg), colony intensity serves as a good proxy for biomass. (Spearman rank correlation r = 0.98, p < 10^-26^). The plot also includes data points from the transition from exponential to stationary phases, which is why the fit becomes less accurate in the top right corner.

**Supplementary Figure 2:**
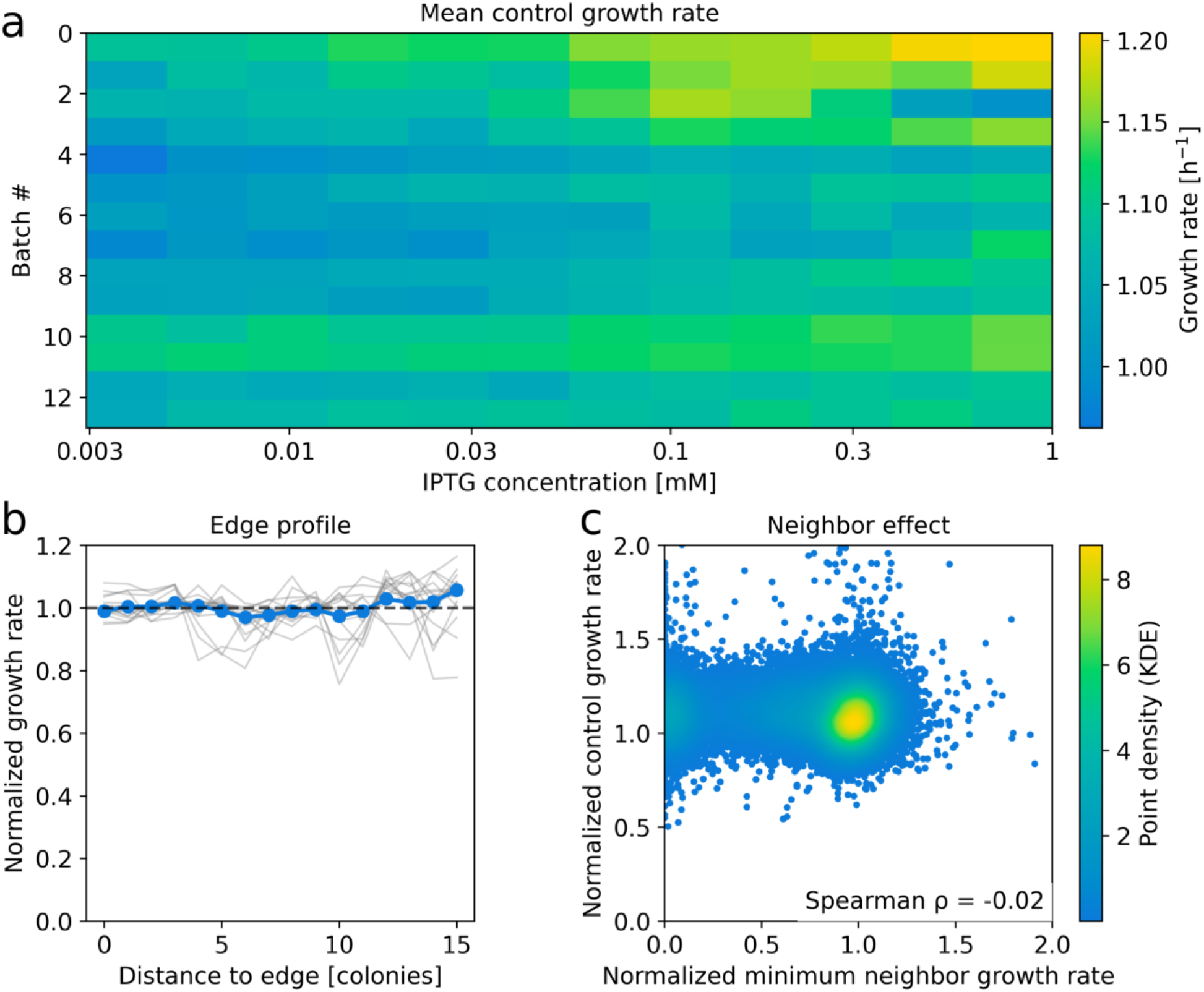
Colony intensity measurements show only weak batch, edge, and neighbor effects, indicating minimal spatial and batch-related biases. **a)** Mean growth rate of control strain for each plate. The control strain carries the ASKA plasmid without an ORF; each plate contains at least 14 colonies of the control strain in different locations (Methods). Rows correspond to batches (ASKA library plates); columns correspond to IPTG concentrations. **b)** Edge profile: control strain growth rates versus distance to the nearest edge. Gray lines show the mean growth rate of all control strain colonies at the same distance from the edge on individual plates; blue line shows the mean across all plates. **c)** Scatter plot of normalized colony growth rate of control strains versus the minimum growth rate among direct neighboring colonies (excluding replicates). Spearman ρ = −0.02, p < 10^-6^.

**Supplementary Figure 3:**
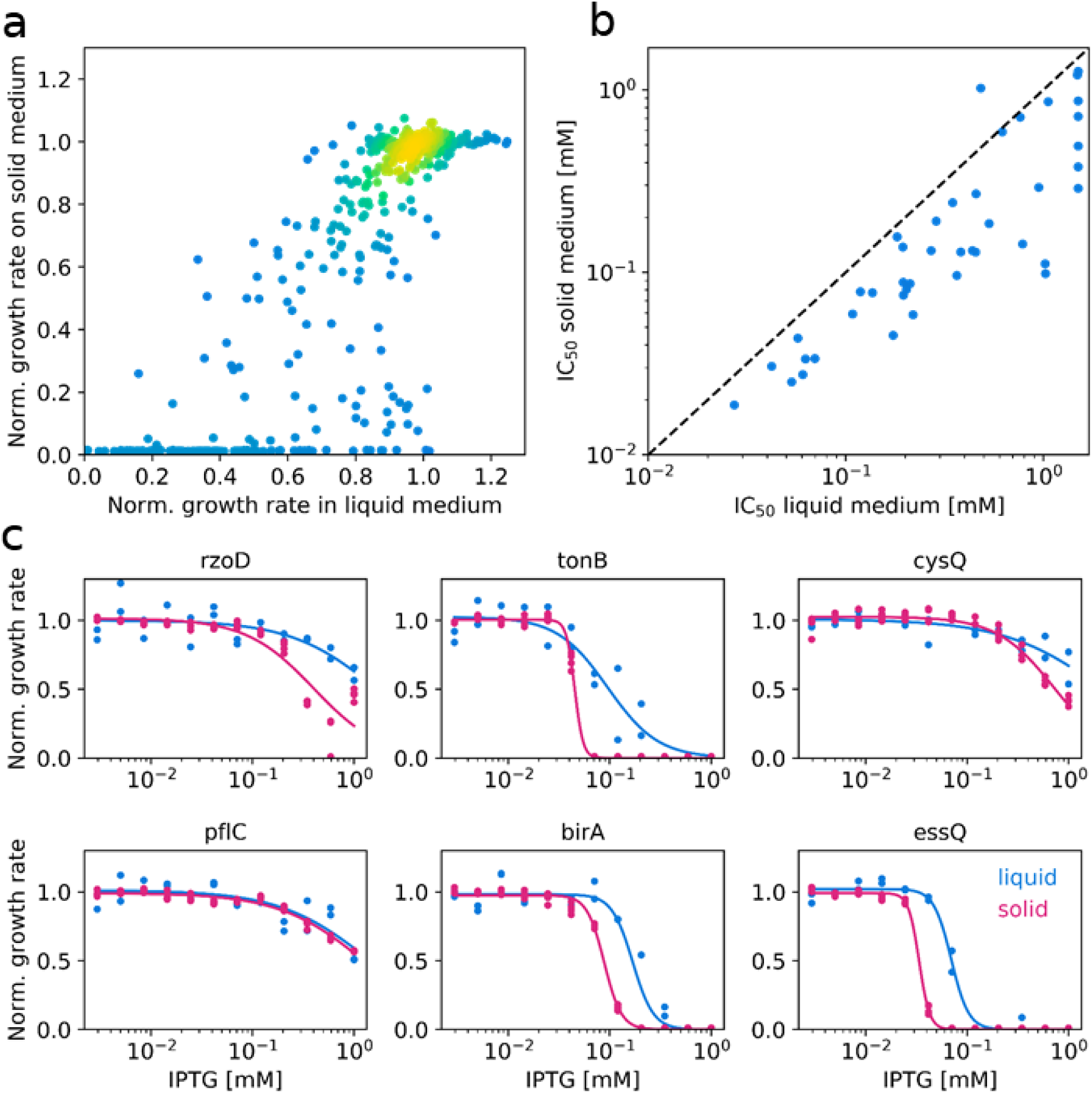
Comparison of growth rates and overexpression dose-response curves measured by colony imaging with those measured in liquid culture. **a)** Scatter plot comparing growth rates of ASKA strains in liquid medium and on solid medium for different IPTG concentrations. Spearman r = 0.79, p < 10^-103^. **b)** Scatter plot comparing IC_50 v_alues from measurements in liquid medium and on solid medium. Spearman r = 0.78, p < 10⁻⁹. The slight systematic offset toward higher IC_50 v_alues in liquid medium is likely due to day-to-day variation in IPTG concentrations or slight IPTG degradation during the preparation of the solid medium (Methods). **c)** Example dose-response curves for six ASKA strains, comparing growth in liquid medium (blue) and on solid medium (magenta). While a few strains (e.g., *tonB*) exhibit divergent behavior that may be linked to the specific function of the overexpressed protein, most strains have similar curve shapes, supporting the robustness of our measurements.

**Supplementary Figure 4:**
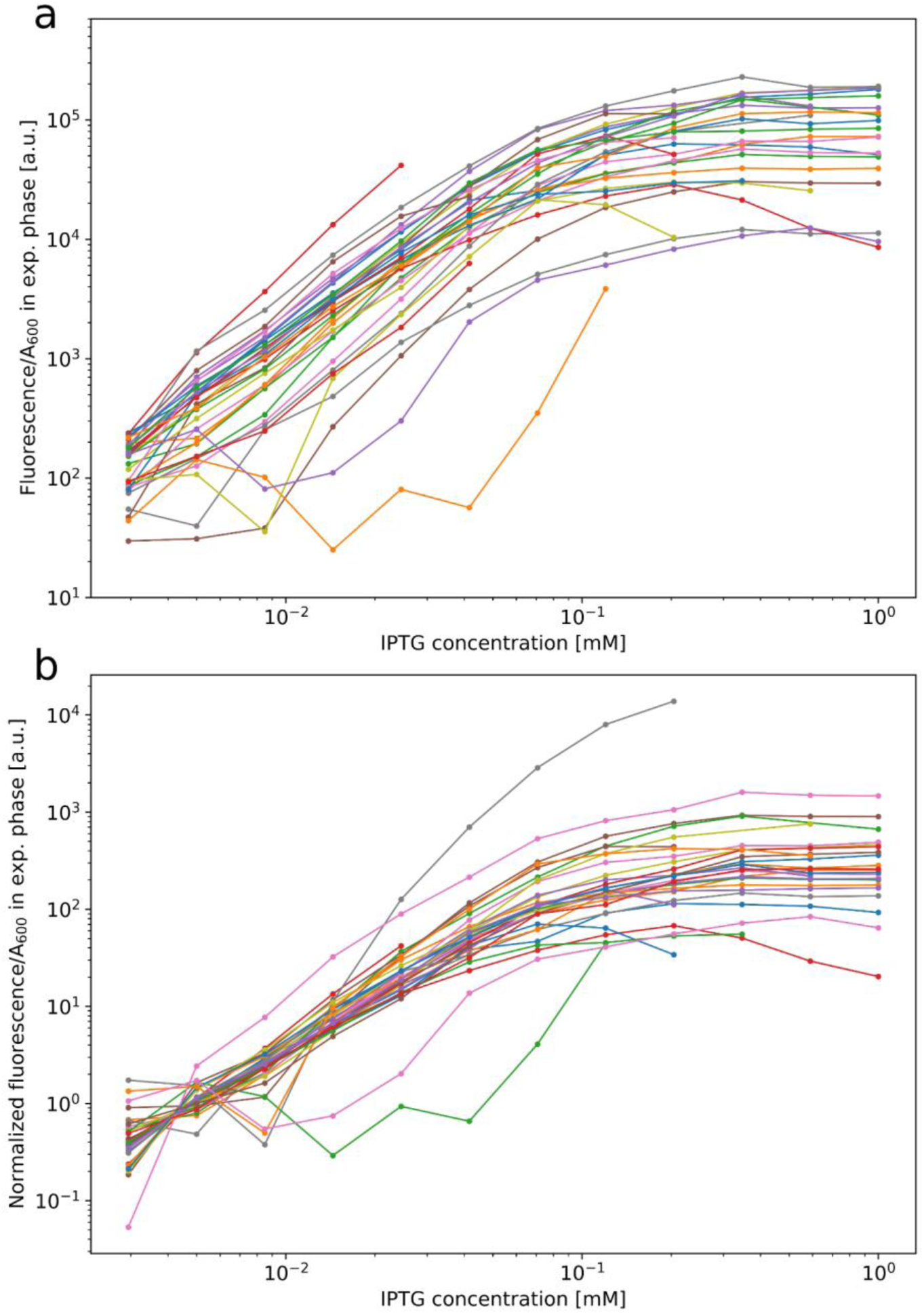
Changing the inducer concentration consistently changes the expression level of different proteins. **a)** Expression levels (GFP/A_600 i_n exponential phase) of 41 ASKA overexpression strains with GFP fusions (Methods) as a function of IPTG concentration. We excluded strains with irregular or discontinuous fluorescence induction curves. Such irregular curves can arise when C-terminal GFP fusions do not reliably report protein abundance because the fusion affects protein folding, or because GFP maturation is impaired. For example, this may occur when the C-terminal end is periplasmic instead of cytosolic (Feilmeier et al., 2000). All data, including gene names, are available in Supplementary Table 2. **b)** Same as **a**, but normalized to the mean of the first two inducer concentrations, which reduces the difference between the individual curves.

**Supplementary Figure 5:**
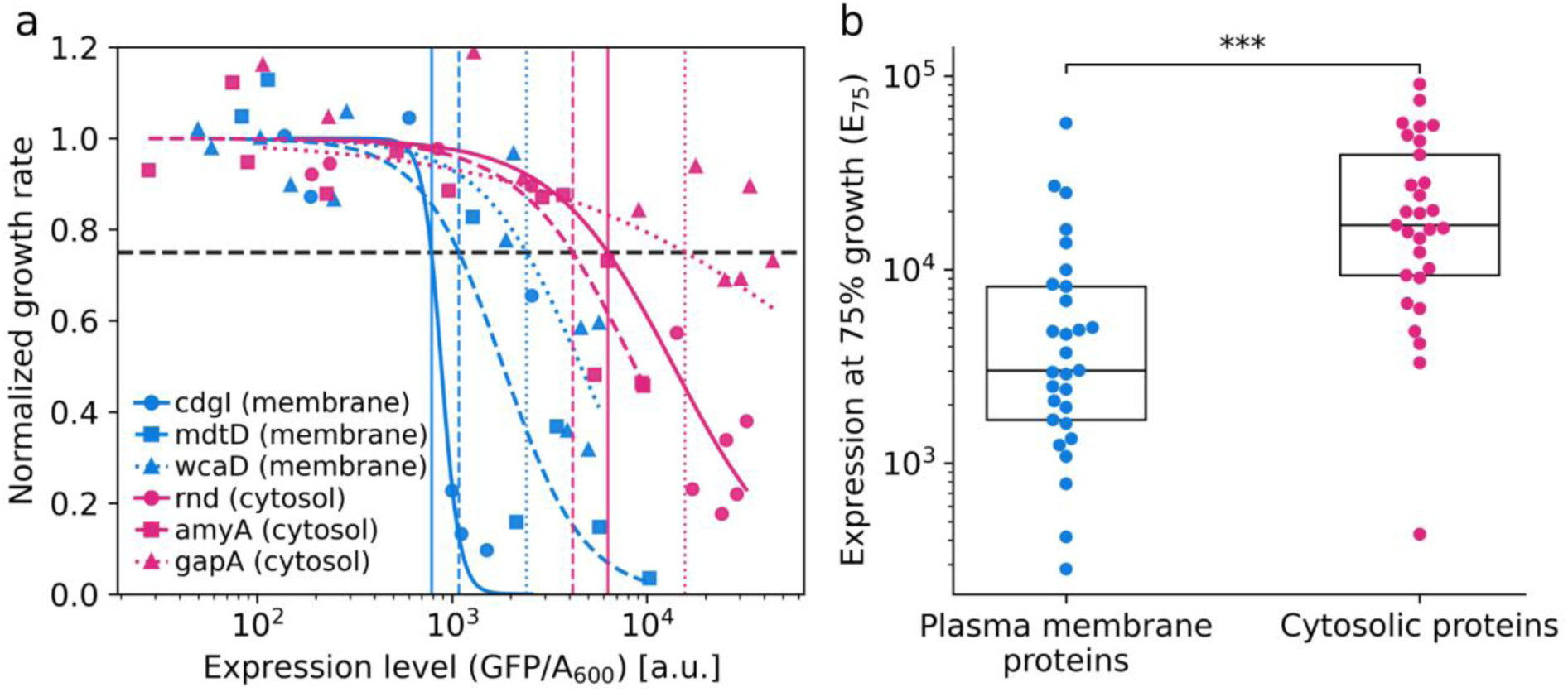
Growth inhibition by membrane protein overexpression occurs at lower expression levels than for cytosolic proteins. **a)** Representative dose-response curves for GFP-tagged ASKA strains, showing normalized growth rate as a function of measured expression level during exponential growth. Expression level was quantified as GFP fluorescence normalized by A_600 (_Methods). Growth rates decreased at lower expression levels for plasma membrane proteins than for cytosolic proteins and the dose-response curves were steeper. Note that the dose-response curves do not collapse onto one universal curve. This is likely due to differences in fitness costs resulting from specific protein functions, variations in the number of TMDs, protein size, and hydrophilicity among other factors. **b)** E75 values, defined as the measured expression level at which normalized growth rate decreases to 75%, for GFP-tagged ASKA strains grouped by protein class. Plasma membrane proteins had significantly lower E75 values than cytosolic proteins, supporting that lower expression levels are sufficient to reduce growth. The difference was significant (*p* < 10^−5^, Mann–Whitney U test). These data are available in Supplementary Table 3.

**Supplementary Figure 6:**
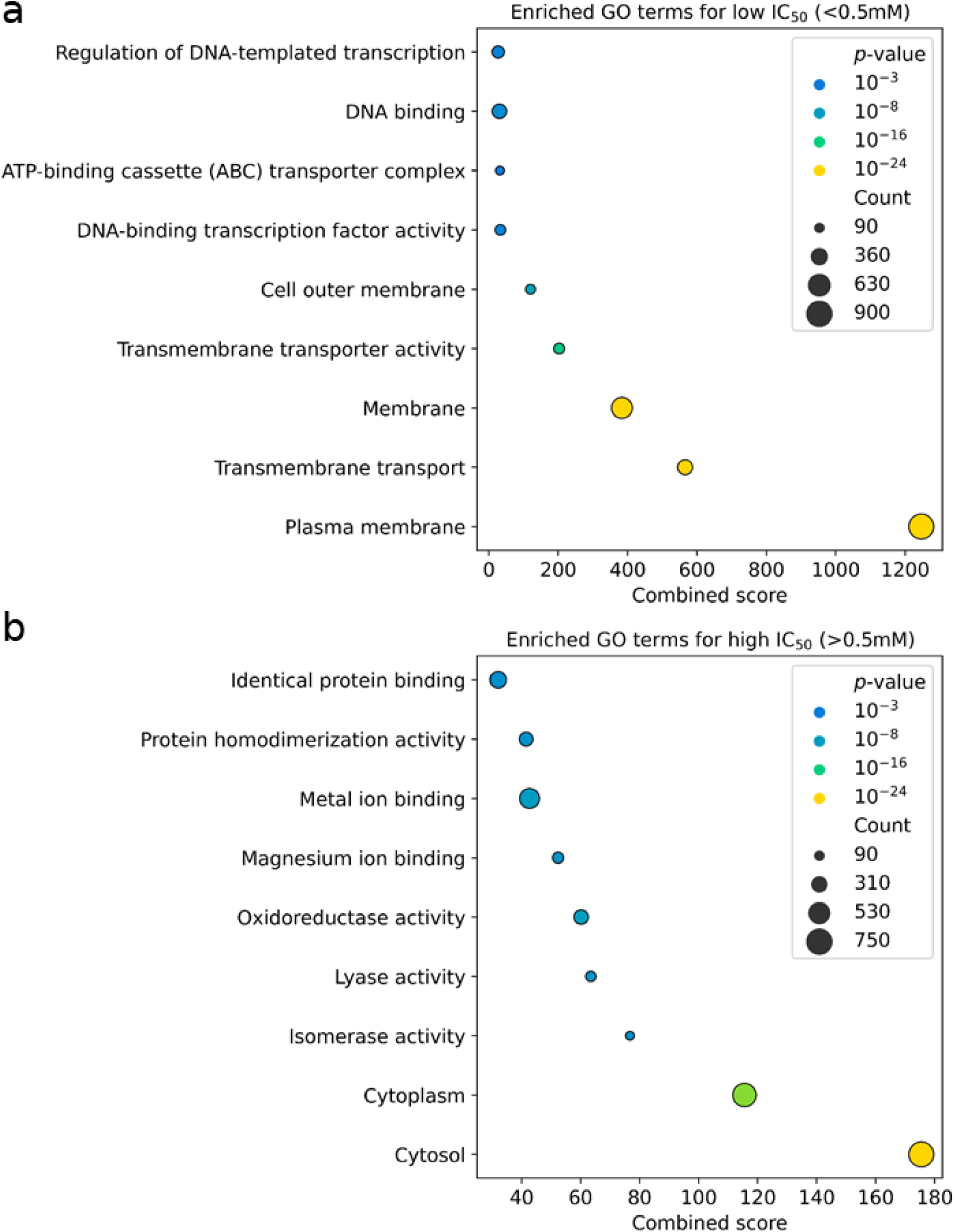
GO enrichment analysis based on fitness cost. **a)** GO terms associated with high fitness cost (IC50 < 0.5 mM). **b)** GO terms associated with low fitness cost (IC50 > 0.5 mM). Only terms with more than 90 genes and *p* < 10^−3^ are shown.

**Supplementary Figure 7:**
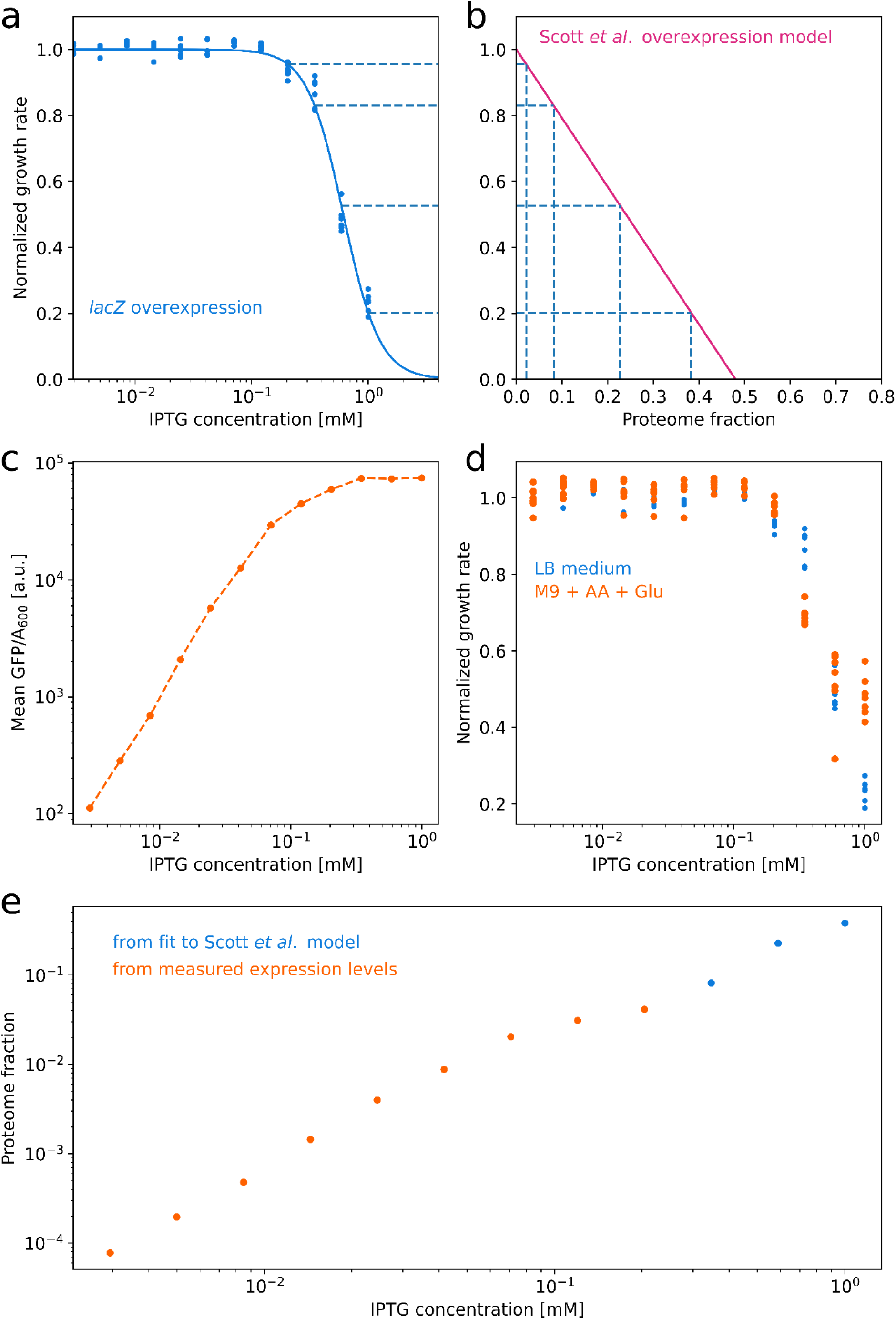
Conversion of inducer concentration to proteome mass fraction. **a)** Measured dose-response curve of the *lacZ* ASKA strain (data points) and Hill function fit (line). **b)** We use the validated sector model of *(Scott et al., 2010)* (magenta line), which was also developed based on *lacZ* overexpression data (Scott et al., 2010), to calculate proteome fractions for the ASKA *lacZ* overexpression under our experimental conditions (dashed black lines). **c)** Mean expression level (GFP signal normalized to A_600,_ which is proportional to optical density) as a function of IPTG concentration for ASKA GFP library strains in M9 medium (Methods). The curve was calculated as the mean of all measured proteins (Supplementary Figure 4). We assume that the expression level of each overexpressed protein follows this trend. **d)** Overexpression dose-response curve of the *lacZ* ASKA strain in LB medium (blue) and the *lacZ*+GFP library strain in M9 medium (orange). The orange curve levels off for high IPTG concentrations, suggesting that maximum induction is reached at lower concentrations in M9 medium than in LB. Consistent with this view, the measured expression levels in **c** saturate at high induction. This difference in the dose-response curves at high IPTG concentrations indicates a difference in the maximum expression levels between the two media. **e)** Inferred proteome mass fractions as a function of IPTG concentration. For the three highest IPTG concentrations, the Scott *et al*. model was used to calculate the proteome fractions. IPTG concentrations of 0.12 mM and 0.20 mM (where the M9 dose-response curve does not yet level off) were used to calculate the conversion factor between GFP/A_600 a_nd proteome fraction. This conversion factor was used to infer proteome fractions for the remaining IPTG concentrations based on the expression level curve measured with GFP/A_600 s_hown in **c**.

**Supplementary Figure 8:**
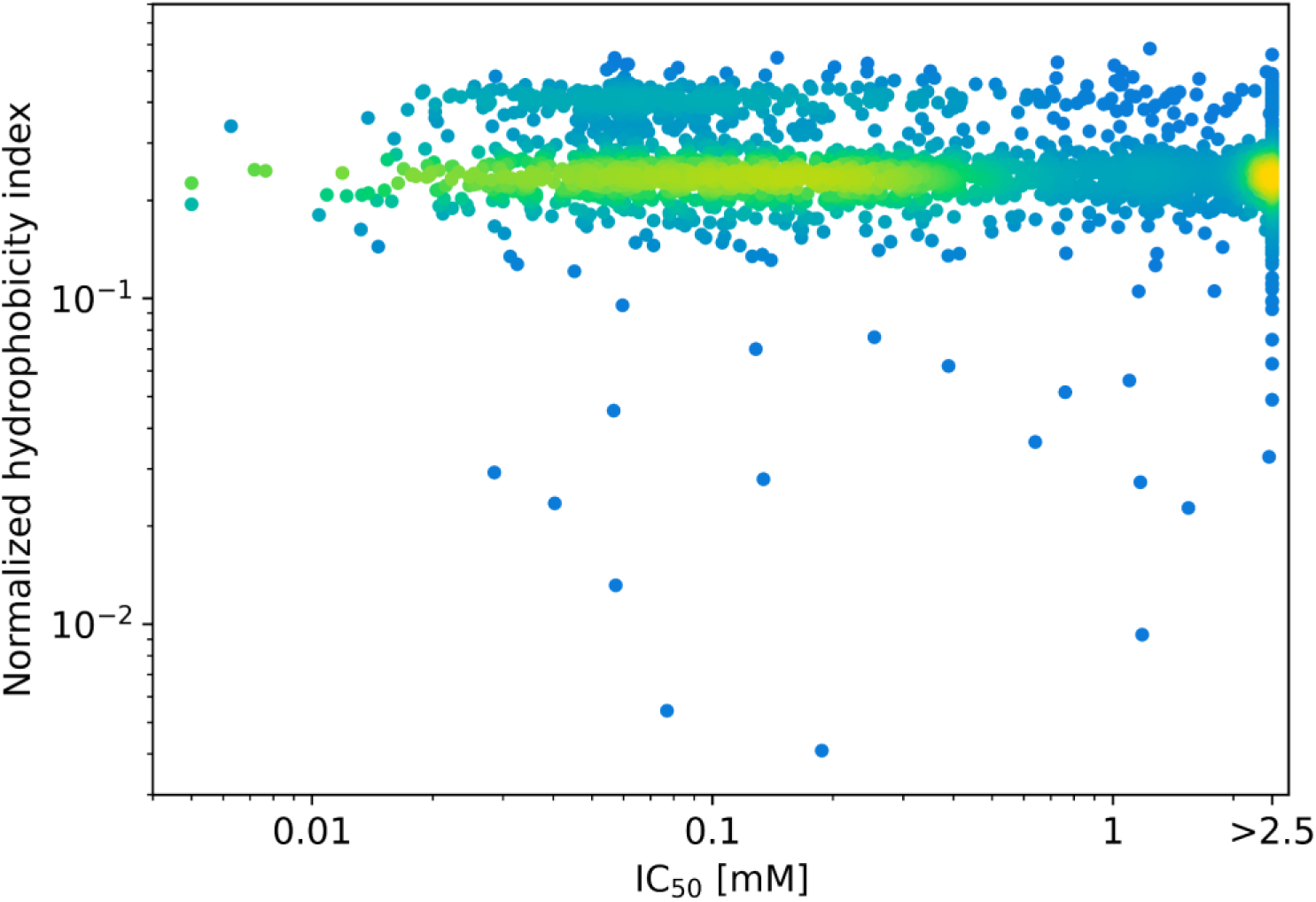
Hydrophobicity and overexpression costs of proteins are weakly correlated. Density scatter plot of hydrophobicity index vs. IC_50 f_or all overexpressed proteins. Proteins with higher hydrophobicity, as measured by the cumulative hydrophobicity index of the amino acids (Methods), tend to have higher fitness costs upon overexpression. Spearman rank correlation *r* = −0.21 (p < 10^−33^).

**Supplementary Figure 9:**
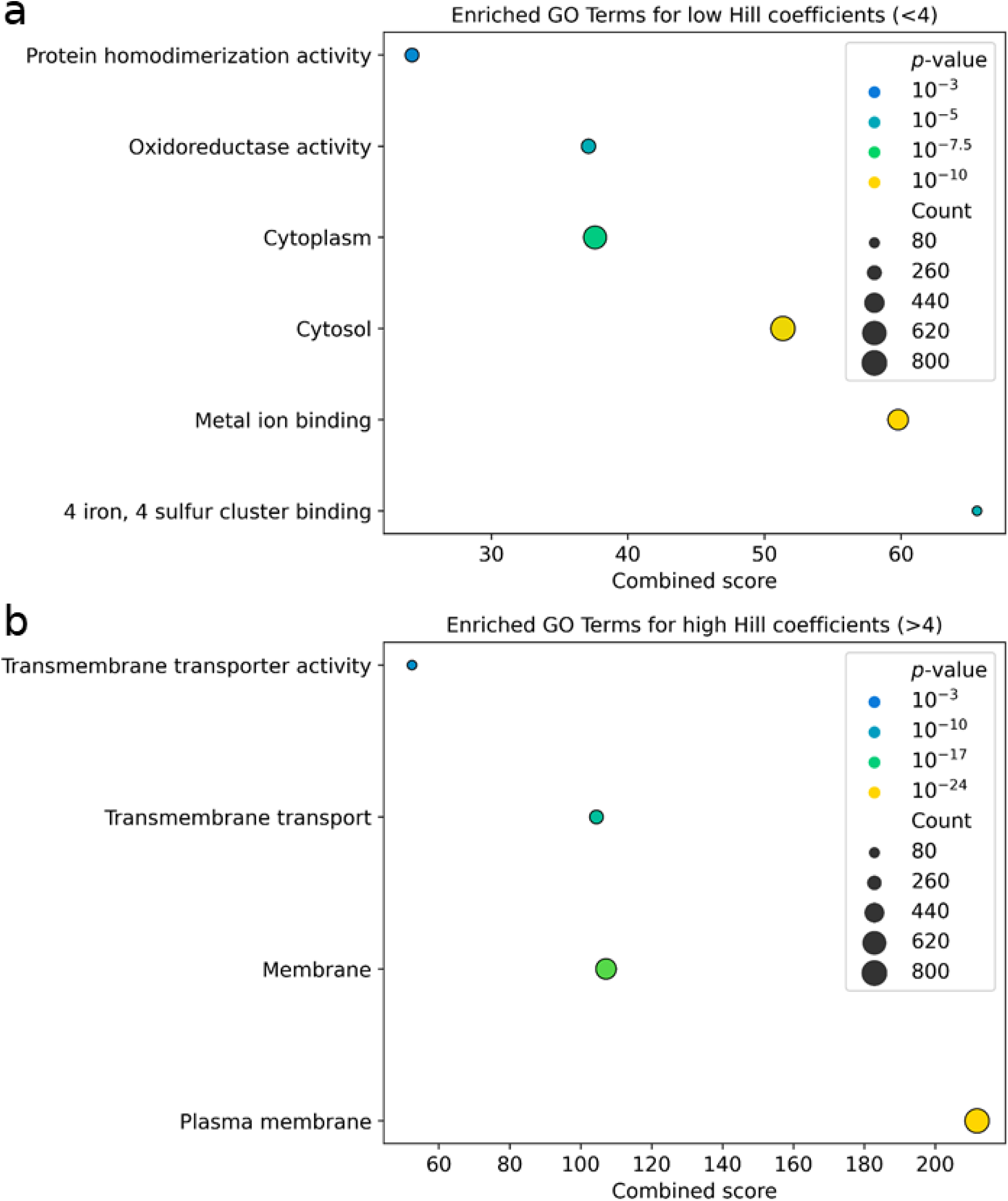
GO enrichment analysis based on overexpression sensitivity. Same as Supplementary Figure 6, but for overexpression sensitivity measured by Hill coefficient (Figure 1d) instead of IC_50_. **a)** GO terms with low overexpression sensitivity (Hill coefficients n < 4). **b)** GO terms with high overexpression sensitivity (Hill coefficients n > 4).

**Supplementary Figure 10:**
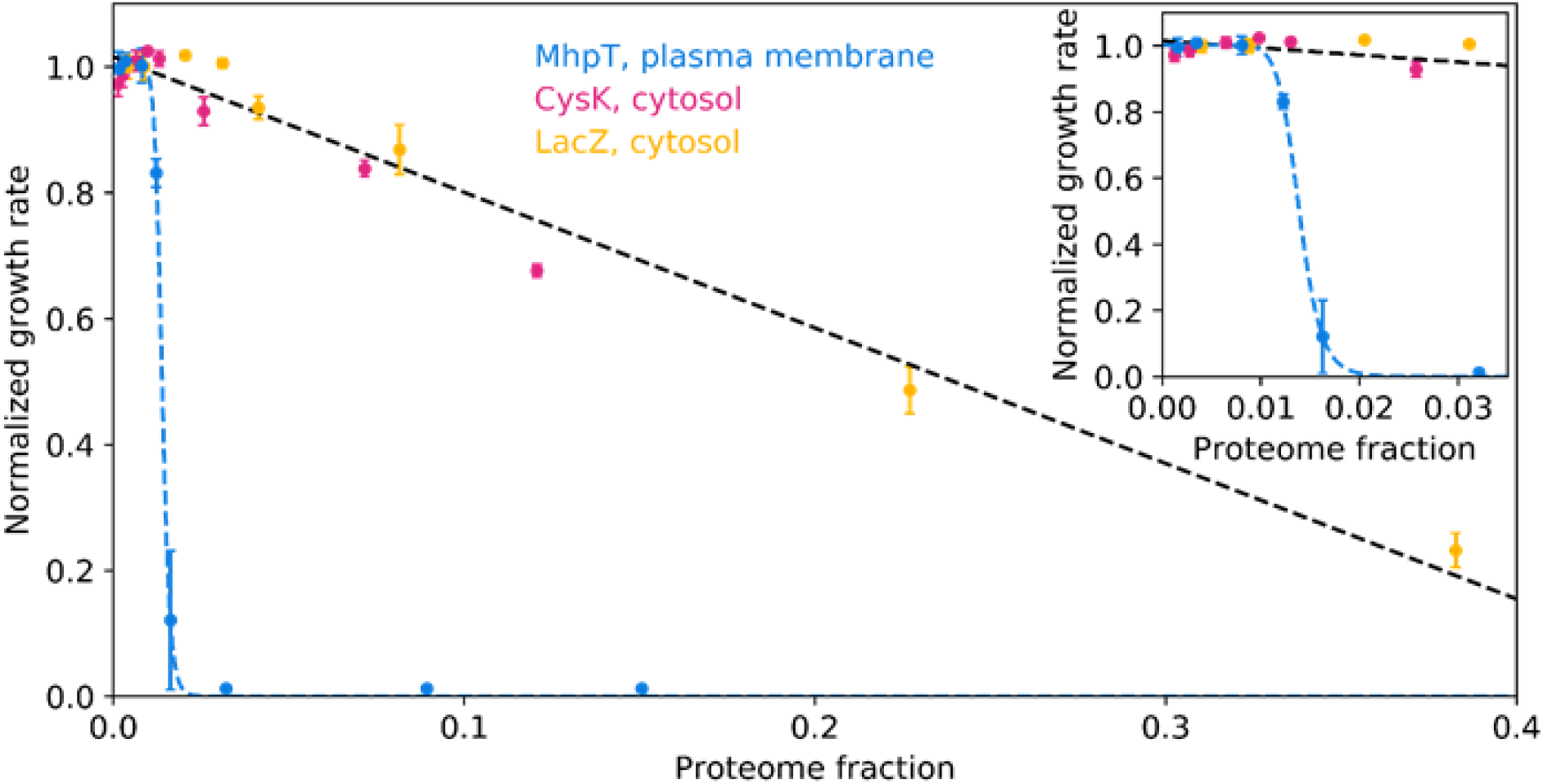
Comparison of growth rate as a function of proteome mass fraction of overexpressed membrane and cytosolic proteins. Overexpression dose-response curves for MhpT (blue), CysK (magenta), and LacZ (yellow) using proteome fractions (Supplementary Figure 7). The proteome fractions were adjusted to account for the different length of MhpT and CysK compared to LacZ. The cytosolic proteins CysK and LacZ show a linear decrease in growth rate (black dashed line), while the membrane protein MhpT shows a distinct non-linear (concave) response (inset: zoom to the lower part of the x-axis).

**Supplementary Figure 11:**
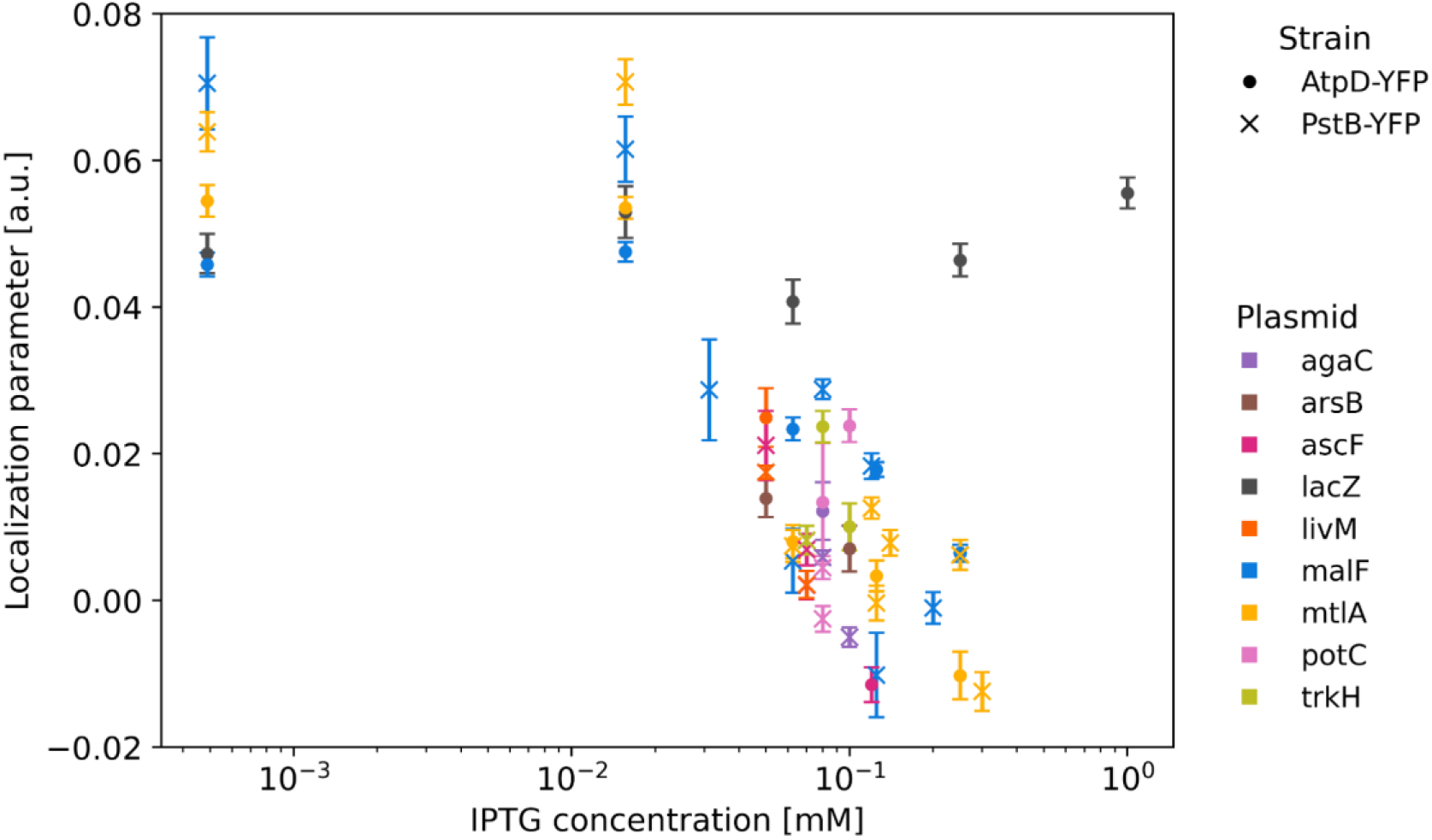
Changes in the localization parameter upon overexpression of different proteins. As Figure 5d, but including additional proteins, as shown in the legend. AtpD-YFP (circles) and PstB-YFP (crosses) were used to quantify membrane localization. The number of TMDs is odd for agaC, arsB and livM; it is even for all other shown membrane proteins.

**Supplementary Figure 12:**
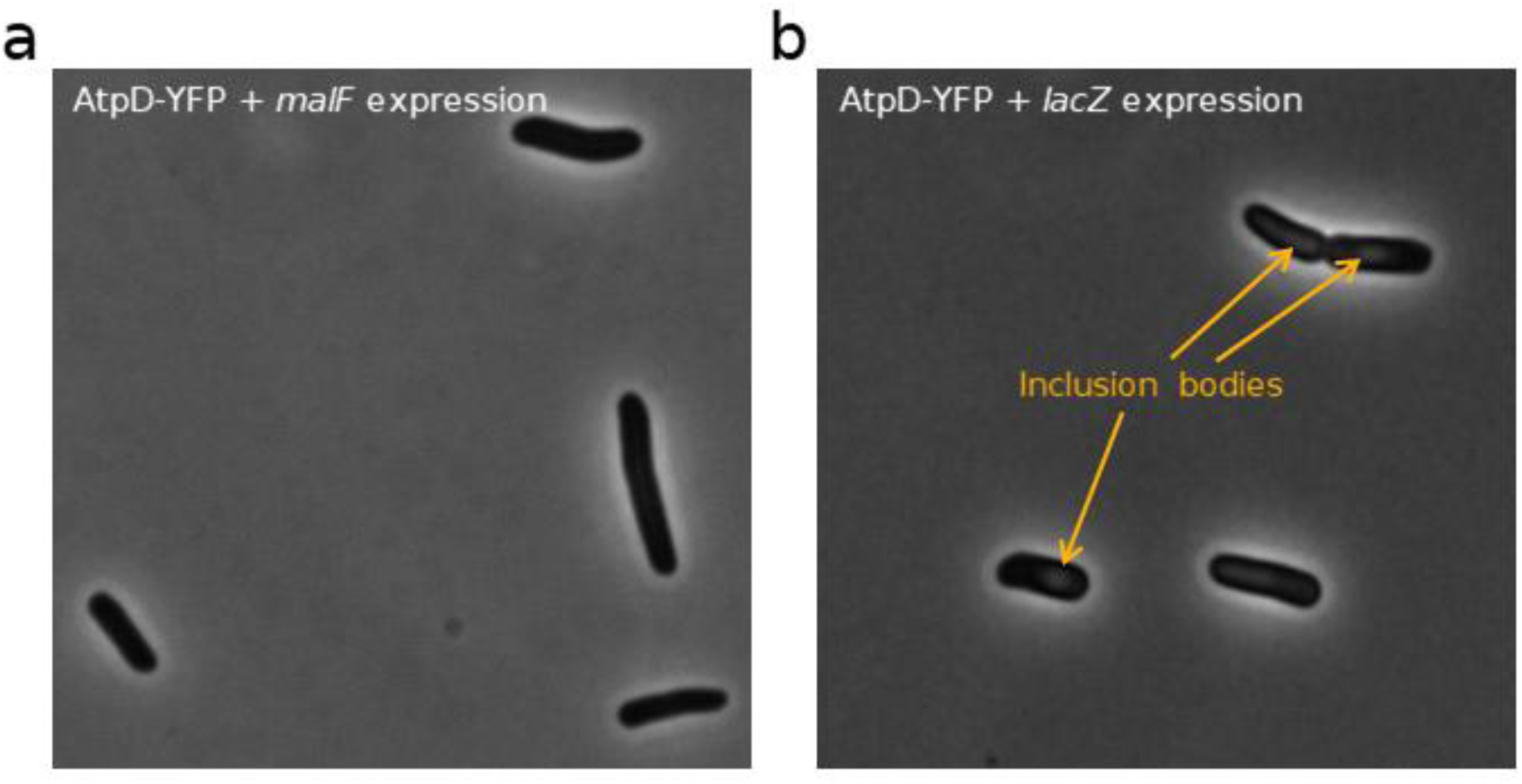
Growth inhibition caused by membrane protein overexpression is not due to protein aggregation or inclusion-body formation. **a)** Phase-contrast image of AtpD-YFP cells overexpressing *malF* at 0.25 mM IPTG, resulting in almost complete growth inhibition. **b)** Same as **a**, but with *lacZ* overexpression at 1 mM IPTG, showing approximately 40% growth inhibition at this inducer concentration.

**Supplementary Figure 13:**
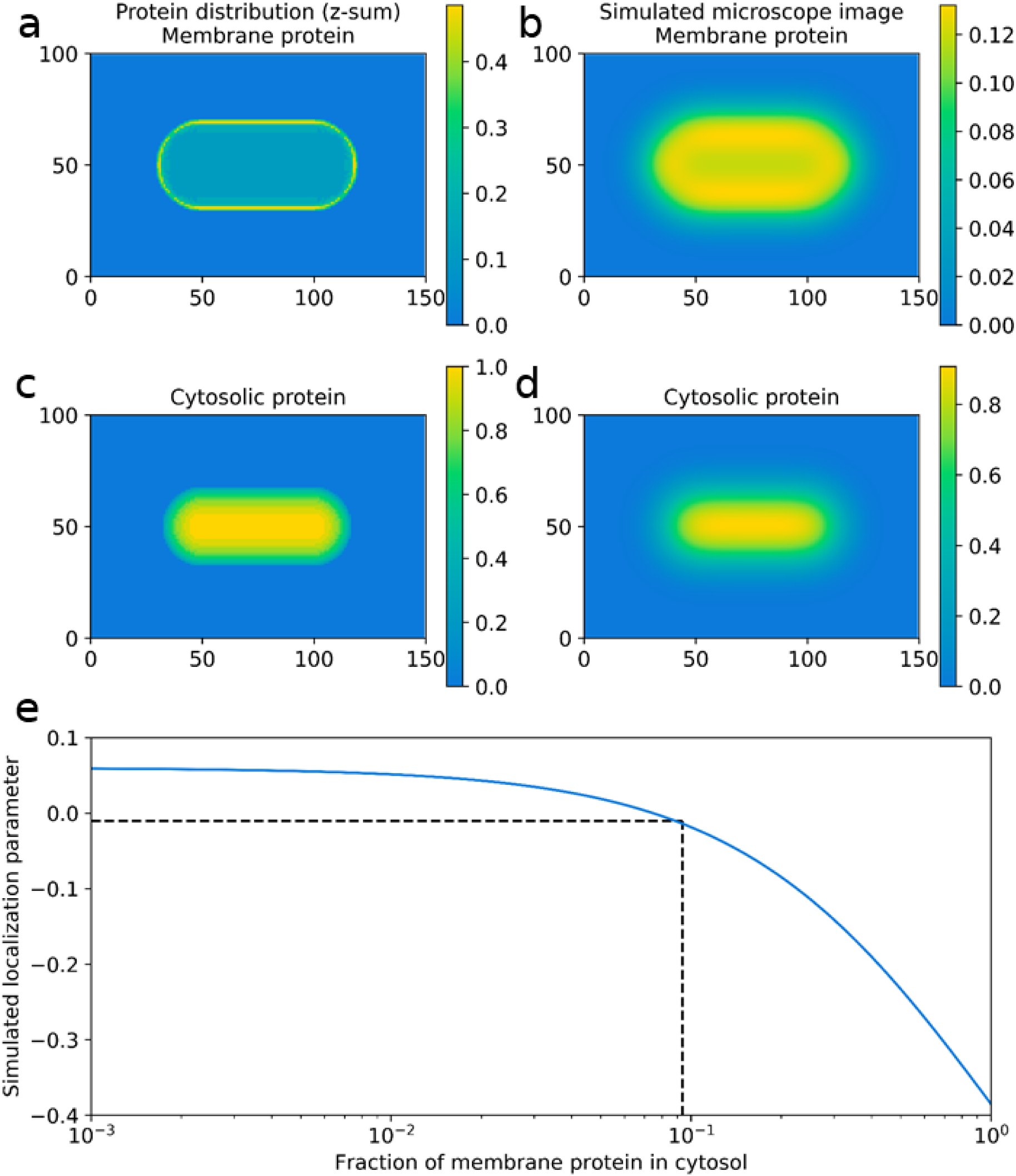
Quantitative interpretation of the localization parameter using a computational model of *E. coli* cell geometry. **a)** Simulated three-dimensional distribution of membrane proteins in a rod-shaped bacterium described as a spherocylinder, shown as a projection along the z-axis. The color represents the protein density per volume normalized to the maximum protein density observed for the cytosolic protein distribution. **b)** Simulated microscope image of the membrane protein distribution in **a**, generated using the point spread function of the imaging setup (Methods). **c)** Simulated three-dimensional distribution of cytosolic proteins in a rod-shaped bacterium, also shown as a projection along the z-axis. **d)** Simulated microscope image of the cytosolic protein distribution in **c**, analogous to **b**. **e)** Simulated localization parameter calculated from composite distributions of membrane and cytosolic proteins with varying ratios (blue line). Experimentally, the growth rate collapses when the localization parameter approaches -0.01, which corresponds to about 9% of the membrane protein being displaced to the cytosol (black dotted line).

**Supplementary Figure 14:**
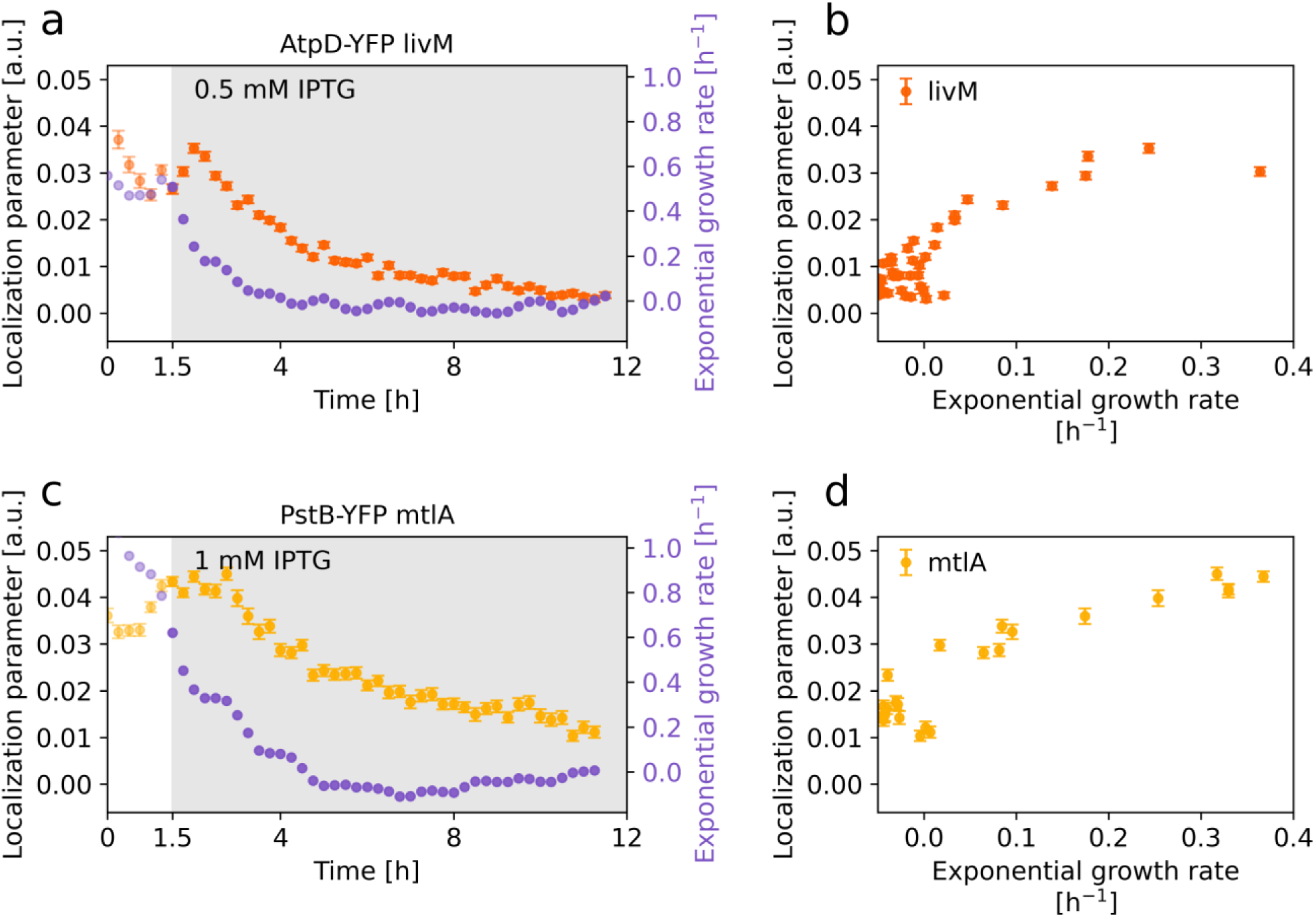
Dynamics of membrane protein displacement for additional membrane proteins. As Figure 6a-c, but for additional overexpressed membrane proteins and an additional membrane localization reporter. **a,b)** Localization parameter for AtpD-YFP and growth rate upon LivM overexpression. Pearson correlation *r* = 0.78, *p* < 10^-8^. **c,d)** Localization parameter for PstB-YFP and growth rate upon MtlA overexpression. Pearson correlation *r* = 0.84, *p* < 10^-11^. Note that, due to different expression levels, the absolute values of the localization parameter differ from those of AtpD-YFP.

**Supplementary Figure 15:**
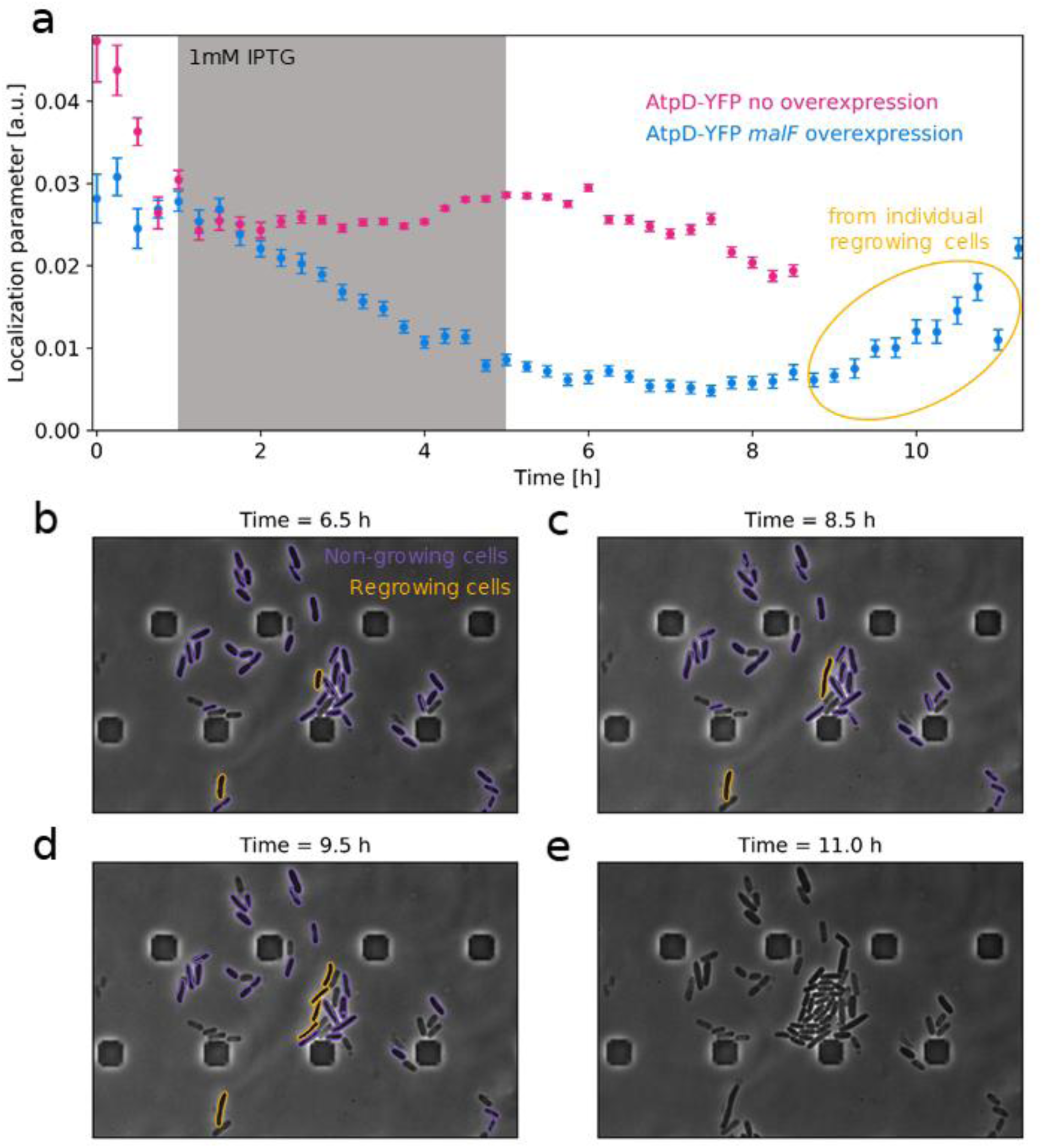
Most bacteria do not resume growth after displacement of endogenous membrane proteins. **a)** Time course of the median localization parameter for single cells tracked in a microfluidic device (Methods). The gray-shaded area indicates the period during which IPTG (at 1 mM) was added to the growth medium. The localization parameter for AtpD-YFP decreases under *malF* overexpression (blue), while it remains approximately constant in a control strain without overexpression (magenta). **b)** Microscopy image from the time-lapse shown in **a**, at 6.5 hours, shortly after IPTG was removed from the medium. Detected bacteria are highlighted in purple and yellow; yellow indicates those bacteria that resume growth within 5 hours. **c-e)** Phase-contrast images from the time-lapse at 8.5, 9.5 and 11 hours. Yellow bacteria gradually resume growth, while purple bacteria remain unchanged.

